# Structural characterization of Posaconazole- parechovirus A3 capsid interaction using molecular simulations and cell infection assays

**DOI:** 10.64898/2026.11.11.765989

**Authors:** Rahil Salehi, Debarpan Dhar, Rangaraj Selvarangan, Ratul Chowdhury

## Abstract

Parechovirus-A3 (PeV-A3) is a major cause of central nervous system infections in neonates, associated with meningoencephalitis and neurodevelopmental delays. Yet there are no specific antivirals against PeV-A3 infection. A newer clade of PeV-A3 has emerged post COVID-19, and our preliminary data shows that infants are not immune to the emerging variants, underscoring the need for a focused therapeutic discovery. Posaconazole (POS) is seen to attenuate PeV-A3 entry and infection, but the mechanistic understanding of drug-capsid interaction remains unknown. To this end, we combined mechanistic interrogation of drug– capsid interactions and conformational dynamics, with experimental infection assays. We identify two POS binding sites: a VP0-VP1 interface in the 2009 isolate (US WI-09), and a VP3-VP1 groove in the 2022 isolate (9C-CMKC) and a laboratory-selected POS-resistant variant (US WI-09 POSA P14). Apo simulations showed congruent trajectories of POS: to the VP0 face in US WI-09, and to the VP3 arm in 9C-CMKC and US WI-09 POSA P14. The relative contribution of VP0 to the drug-capsid interaction decreased from 86% in US WI-09 to zero in both 9C-CMKC and US WI-09 POSA P14, indicating a redistribution of binding interactions across capsid subunits. Cysteine or arginine substitutions to VP1 Tyr224 abolished the Tyr224-VP3 anchor. Functionally, 0.3 µM POS inhibited replication of US WI-09 and 9C-CMKC in Vero-P cells but only US WI-09 in receptor-overexpressing Vero-MYADM cells. These findings suggest that POS efficacy depends on binding-site localization near the MYADM-engaging VP0 interface, revealing how emerging PeV-A3 variants may retain drug binding while evading antiviral inhibition.

**Importance:** PeV-A3 causes neonatal sepsis and central nervous system infection with long-term neurodevelopmental sequelae, and clinical care remains supportive. Posaconazole, an antifungal azole compound is reported to inhibit PeV-A3, yet their capsid target was undefined, and isolates circulating since 2022 form a distinct clade of unknown susceptibility. We show that posaconazole (POS) acts by occupying an uncleaved VP0-VP1 interface adjacent to a known resistance position, and that both laboratory-selected resistance and natural clade divergence relocate the drug to a VP3 pocket that binds it as tightly while leaving the entry-critical VP0 face free. A favorable computed affinity at a non-productive site is therefore indistinguishable from efficacy by binding metrics alone. This reframes how capsid-directed antivirals should be screened, identifying the VP0 surface as the next-generation target for PeV-A3 inhibition, and supplies a workflow that transfers directly to other picornaviruses.

## 1. INTRODUCTION

Parechovirus A (PeV-A) is a common childhood pathogen in the *Picornaviridae* family and can cause severe illness in infants (1). There are 19 reported PeV-A genotypes, and most infections are mild, with gastrointestinal or respiratory symptoms. However, infection with the PeV-A3 genotype is associated with severe diseases in infants under three months of age. Parechovirus-A3 (PeV-A3) is among the most frequently detected picornaviruses in cerebrospinal fluid (CSF) or blood from infants suspected of sepsis or central nervous system (CNS) infections (2–4) with incidences following a biennial pattern (5, 6). Clinical presentations of infected infants include poor feeding, lethargy, fever, rash, irritability, with severe cases leading to meningoencephalitis (1, 7). Severe cases of PeV-A3 infection have shown white matter abnormalities during initial hospitalization (8). Furthermore, long-term neurological sequelae and neurodevelopmental delay have also been associated with PeV-A3 infections (9–11). Some of the observations include motor delays, cognitive deficits, behavioral issues, speech/language delays, and visual problems (11, 12). No specific antiviral therapy is available for PeV-A3 infection. Care is supportive, and intravenous immunoglobulin is sometimes used. Evidence from previous literature indicates that selected antifungal azoles, including itraconazole (ITZ) and posaconazole (POS), inhibit PeV-A3 *in vitro* by binding with the viral capsid, and inhibiting the internalization of PeV-A3 into host cell (1, 13, 14). However, the precise POS binding site(s) on the PeV-A3 capsid remains undetermined. In addition, newer PeV-A3 clinical isolates from 2022 are significantly divergent in amino acid sequences from isolates of previous years, forming a separate clade (15), that could likely inhibit POS binding with the capsid. A structure-guided mechanistic interrogation of PeV-A remains understudied; that hinders a rational design of antiviral against PeV-A3.

All-atom molecular dynamics (MD) simulations can help resolve capsid motions under physiological solvent conditions and can detect small changes in stability, flexibility, and allosteric coupling after drug binding that are not obvious in static models, such as crystal and cryo-EM structures(16, 17). When an experimental capsid-drug complex is unavailable, docking offers a possible starting point, placing a ligand into the candidate pocket prior to MD refinement (17). Classical docking is fast but often imprecise for reliable affinity ranking, whereas free energy perturbation (FEP methods) offers greater accuracy at substantially higher computational cost (18). Deep learning models for biomolecular complexes have narrowed the gap between static structure prediction and ligand ranking. Boltz-1 reached comparable performance on protein-protein and protein-ligand benchmarks and was released openly(19). Boltz-2 builds on that framework with geometric deep learning and diffusion-based all-atom prediction, returning complex structures together with confidence metrics such as pLDDT and interface scores, and providing an alternative to classical docking for pose generation (18, 20). Boltz-2 predicts all-atom complex structure and small-molecule binding affinity in one pass and builds on the co-folding approach of Boltz-1(19) and AlphaFold3(21). Critically, Boltz-2 also reports an affinity estimate, offering a route to ligand ranking that is significantly faster than atomistic free-energy methods while approaching their accuracy on established benchmarks (18, 22, 23). These features make Boltz-2 an ideal starting tool to position POS on PeV-A3 capsid models derived from protein database (PDB) and compare predicted poses and affinities across different clinical isolates and resistant mutants prior to MD refinement. MD refinement then relaxes the predicted complex, yields a trajectory-averaged estimate of binding affinity, and reports whether the ligand remains engaged or moving beyond a single docking/co-folding score, as in our prior PRRSV docking and relaxation-based binding-energy ranking (24, 25).

Capsids are favored antiviral targets as they regulate important steps during viral infection, and small-molecule drugs that bind to or interfere with functioning of the viral capsid can block viral replication (16, 17). In enteroviruses, classical capsid inhibitors bind the hydrophobic VP1 pocket normally occupied by a lipid pocket factor. By replacing this factor, these compounds stabilize the virion, increase the energetic barrier to receptor- or thermally induced uncoating, and thereby prevent the conformational transitions required for genome release (26, 27). A cryo-EM structure of an enterovirus, coxsackievirus A10, supported virtual screening of about four million compounds into that VP1 pocket and yielded inhibitors that were confirmed using *in vitro* cell-based assays (28). In other studies, a combination of cryo-EM, resistance genetics, and MD refinement help identify a second drug binding site at a VP1-VP3 interprotomer interface, that prevents the expansion needed for RNA release (27, 29). PeV-A3 are different as their atomic structure lacks the hydrophobic VP1 pocket, rendering the pocket-binding anti-picornavirus compounds ineffective against PeV-A3 infection (30).

To define the posaconazole binding site on PeV-A3, Boltz-2 docking of two-protomer capsid models was followed with all-atom MD refinement of POS complexes comparing two PeV-A3 isolates: US WI-09 (isolate from 2009) and 9C-CMKC (isolate from 2022). Pocket contacts, cohesion, and molecular mechanics/Poisson-Boltzmann surface area (MM/PBSA) decomposition were used to compare occupancy, chemistry, and binding energies (enthalpy) between isolates. Finally, replication of PeV-A3 in the presence of POS was studied to validate the impact of differential drug-capsid interaction between isolates on overall viral replication. Together, this work provides the first integrated mechanistic deep dive structural, dynamic, and functional characterization of POS binding on PeV-A3 capsid. To our knowledge, no published study has docked POS onto PeV-A3, run MD refinement of a POS-capsid complex, and/or compared predicted POS engagement across PeV-A3 clinical isolates of two different clades.

## 2. METHODS

### 2.1. Viral cryo-EM structure and PeV-A3 clinical isolate sequences

The cryo-EM structure of PeV-A3 virion bound to the neutralizing Fab AT12-015 (PDB 6GV4, 2.8 Å) was used as the starting model (31). The Fab heavy and light chains and the ordered genomic RNA were removed, leaving the capsid proteins VP0, VP1, and VP3. POS binds an interface between subunits rather than inside a single protein, so the modeling unit was built from two adjacent capsid protomers. Each protomer contributes one copy of VP0, VP1, and VP3, so the two-protomer assembly has six protein chains: A, B, and C for the first protomer and D, E, and F for the second. Working with two protomers instead of an isolated subunit keeps the inter-protomer contacts that form the drug pocket, following the reasoning used in other picornavirus and capsid drug-binding simulations (17, 32). Three PeV-A3 capsid sequences were used, namely, US WI-09 (MN781635.1; 2009 isolate), 9C-CMKC (PV290892; 2022 isolate), and US WI-09 POSA P14 (MN781636.1), which is a POS resistant variant carrying the four capsid substitutions, VP0 K66R, VP0 A124T, VP3 N88S, and VP1 Y224C as described previously (14, 33). The PeV-A3 capsid sequences were modeled on a two-protomer unit.

### 2.2. Docking to construct drug-capsid complex and affinity prediction

Preliminary POS-capsid complexes and binding-affinity estimates were generated with Boltz-2 (18) for PeV-A3 US WI-09, 9C-CMKC, and US WI-09 POSA P14^19^ For each strain, the input listed the six protein chains of the two-protomer unit and POS as an isomeric SMILES string, with POS set as the affinity binder, so the run returned both a pose and an affinity value. Within Boltz-2 we used the built-in MSA server for alignment generation and the diffusion potentials option. Structure sampling used one diffusion sample with 25 sampling steps, and affinity prediction used five diffusion samples with 200 sampling steps. To check pose reproducibility, each isolate was predicted across 25 independent random seeds, and structures were written in mmCIF format. The complex used to seed molecular dynamics (MD) was taken from these runs.

### 2.3. Ligand (drug) and protein (capsid) parameterization for Molecular Dynamics (MD)

POS was parameterized with the general AMBER force field GAFF2(34). Atomic charges were assigned with the AM1-BCC method(35) for a neutral ligand using Antechamber, missing parameters were filled with parmchk2, and the topology was converted to GROMACS format with ACPYPE(36). The protein was described with the AMBER99SB-ILDN force field (37), and the two-protomer topology was built with pdb2gmx after stripping the ligand heteroatoms. A single POS molecule was then merged with the six-chain protein to form each starting complex. The same parameterization was applied to PeV-A3 US WI-09, 9C-CMKC, and US WI-09 POSA P14.

### 2.4. Structure refinement and scoring drug-capsid interactions with MD

All-atom simulations were run in GROMACS 2023 (38). Each two-protomer complex was centered in a dodecahedral box with a 1.2 nm minimum distance to the box edge, solvated with TIP3P water (39), and neutralized with Na+ and Cl− ions added to 0.15 M. A two-protomer system was chosen as the sufficient capsid definition since it captures all possible inter-protomer interfaces. The system was energy-minimized by steepest descent to a maximum force below 1000 kJ mol⁻¹ nm⁻¹. Equilibration ran in two 100 ps stages with harmonic restraints on the protein heavy atoms. The first stage was NVT at 300 K with the velocity-rescale thermostat (40) and initial velocities drawn from a Maxwell-Boltzmann distribution. The second stage was NPT at 300 K and 1 bar using the velocity-rescale thermostat and the Berendsen barostat (41). Temperature was coupled separately for the protein and non-protein groups in both stages. In line with standard practice, the system was first equilibrated at constant volume (NVT) to stabilize the temperature, since simultaneous velocity generation and box rescaling can destabilize a system far from equilibrium. Pressure coupling was then applied (NPT) to relax the solvent density to the constant-pressure conditions used in production (42, 43). Protein heavy-atom restraints were maintained throughout, allowing solvent reorganization without distortion of the capsid (42).

Production runs were 500 ns per system with no restraints. They used the leap-frog integrator with a 2 fs time step, the velocity-rescale thermostat at 300 K, and the Parrinello-Rahman barostat at 1 bar(44). Bonds to hydrogen were constrained with LINCS (45), long-range electrostatics used particle-mesh Ewald with a 0.16 nm grid (46, 47), and short-range electrostatic and van der Waals interactions used a 1.0 nm cutoff with a long-range dispersion correction. One-ligand starting states were simulated for every system. In the holo state, POS started in the Boltz-2 predicted pocket. In the displaced state, POS was translated about 10 Å away from the holo pocket and then simulated without restraints. The displaced runs are rebinding controls rather than ligand-free apo simulations: the ligand stays in the system but begins roughly 10 Å from the site, so any later contact reflects the local accessibility of the pocket instead of the pre-placed pose. This gave six independent 500 ns trajectories, each on the two-protomer assembly with one POS molecule: ‘wild-type’ US WI-09, 9C-CMKC, and US WI-09 POSA P14 each in the holo and the displaced state.

Global stability was measured by backbone and protein RMSD, POS RMSD after fitting on the protein, per-residue RMSF, and radius of gyration. Ligand engagement was followed by the minimum protein-POS distance, the protein-POS center-of-mass distance, and protein-POS hydrogen bonds. Solvent-accessible surface area, water radial distribution around the protein and POS, mean-squared displacement, and energy terms (temperature, pressure, potential, volume) were also recorded. Conformational sampling was summarized by linkage clustering at a 0.12 nm RMSD cutoff and by principal-component analysis on the backbone. The six trajectories were compared across isolates in the holo and displaced states and as paired holo versus displaced runs within each isolate.

### 2.5. Pocket and mutation contact-network analysis

Residue contact networks around the POS binding site were built from the MD trajectories with MDAnalysis (48, 49) and NetworkX (50). A residue was considered in contact with POS when any of its heavy atoms lay within 4 Å of a POS heavy atom, the criterion used to define protein-ligand contact frequency in MD trajectories (51). Pocket residues were connected in the network when their side-chain heavy-atom centroids (Cα for glycine) lay within 6 Å, within the 4-6 Å range evaluated for side-chain centroid networks derived from MD ensembles (52). The local environment of the site was defined as all residues with a heavy atom within 12 Å of POS, a ligand-centered cutoff previously used to delimit the binding-site region (53). Networks were computed with a stride of five frames, and shortest-path searches between substitution sites and the pocket were capped at six hops. The four resistance positions in US WI-09 POSA P14 (VP0 K66R, VP0 A124T, VP3 N88S, and VP1 Y224C) were tracked in every system at the equivalent positions to test whether each substitution contacts the pocket directly, connects to it through a persistent contact path, or stays decoupled during the simulation.

### 2.6 Binding free energy (MM/PBSA)

Binding energetics were estimated with gmx_MMPBSA (54), which runs end-state Molecular Mechanics Poisson-Boltzmann Surface Area calculations on GROMACS trajectories through the MMPBSA.py engine (55). Calculations used the single-trajectory protocol on the 500 ns production runs for all six systems, US WI-09, 9C-CMKC and US WI-09 POSA P14 in both the holo and displaced states. The receptor was defined as the union of pocket residues around POS and the ligand as POS, and about 500 frames were sampled at a fixed interval across each trajectory. The polar solvation term was computed with the Poisson-Boltzmann model at an internal dielectric of 1, and the nonpolar term was taken from the solvent-accessible surface area. Per-residue decomposition was carried out for residues within 4 Å of POS. The MM/PBSA step was run with GROMACS 2024.4 and AmberTools 22. Conformational entropy was estimated with the interaction-entropy method (56) over the final quarter of the sampled frames. Because interaction-entropy values fluctuated strongly for these large, flexible interfaces, the binding enthalpy (ΔH) is reported as the main comparative metric and the full free energy is read with caution, following the same practice used in our multi-epitope vaccine work, where gmx_MMPBSA enthalpies rather than entropy-corrected free energies were used to rank related complexes (57).

### 2.7 Cells, viruses and primers

The Vero-Polio (Vero-P) cell line, as previously described(14), was provided by collaborators at the Centers for Disease Control and Prevention, Atlanta, USA. Vero-MYADM cell line, as described previously(58) was a generous gift from the Jan Carette Lab at Standford Medicine, USA and Matthew Vogt Lab at University of North Carolina, Chapel Hill. Vero-P and Vero-MYADM cells were maintained in Eagle’s Minimum Essential Medium (EMEM) (ATCC, Catalog #30-2003) supplemented with 10% Fetal Bovine Serum (Catalog #16140071, Thermo Fisher Scientific) and 1% Penicillin–Streptomycin (Catalog #15070063, Gibco).

The two PeV-A3 clinical isolates, US WI-09 (MN781635.1; 2009 isolate) and 9C-CMKC (PV290892; 2022 isolate) were provided by the Centers for Disease Control and Prevention, Atlanta, USA. Upon receipt, these viral isolates were passaged twice in Vero-P cells. Total nucleic acids were extracted using the KingFisher APEX MagMAX^TM^ Viral/Pathogen Nucleic Acid Isolation Kit (Thermo Fisher Scientific, Cat # A48310) and viral titer (genome copies/ml) was estimated using a single step RT PCR using qScript XLT 1-Step RT-PCR Kit (Quantabio, Cat# 95134-500). The primer and probe sequences used for viral titration are AN344 (5′-GGC CCC WGR TCA GAT CCA YAG T -3′), (AN345; 5’-GTA ACA SWW GCC TCT GGG SCC AAA AG -3′) and AN818 [5′-(Cy5)- CCT RYG GGT ACC TYC WGG GCA TCC TTC-BHQ2 -3′]. The thermocycling conditions are cDNA synthesis stage (50°C, 30 min followed by 95°C, 1 min) and PCR stage of 45 cycles (95°C, 10 sec followed by 60°C, 45 sec) on ABI7500 Fast Real-Time PCR instrument (Thermo Fisher Scientific, Cat # 4351106).

### 2.8. Time course replication assay

Vero-p and Vero-MYADM cells were plated in 24 well culture plates at a seeding density of 1×10^5^ cells/ well. After overnight incubation, assuming cells have doubled, respective wells were either mock infected or infected with PeV-A3 USWI-09 and PeV-A3 9C-CMKC at an MOI of 1 (2×10^5^ genome copies/ml), with or without POS at a concentration of 0.3uM as described previously (14). Five days post infection, cells were freeze-thawed three times, and total nucleic acid was isolated from whole cell lysate, and viral titration was done as previously described.

### 2.9. Data availability

All data supporting the conclusions of this study are presented in the article and its supplemental material. Coordinate files for the three Boltz-2 co-folded capsid–posaconazole complexes (US WI-09, US WI-09 POSA P14 and 9C-CMKC) are provided as supplemental material. The PeV-A3 cryo-EM structure used as the modelling template is available in the Protein Data Bank under accession 6GV4 (https://www.rcsb.org/structure/6GV4).

## 3. RESULT

### 3.1. POS occupies VP0-VP1 binding site in US WI-09 but a very different VP3-VP1 site in 9C-CMKC

A single asymmetric unit of PeV-A3 from PDB 6GV4, as the starting template (31) revealed intra-protomer pockets, but as a weak place to look for drug sites between neighboring units. Boltz-2 co-folding was therefore run on a two-protomer capsid unit (six chains: 2×VP0, 2×VP1, and 2×VP3) with POS (18). The aim was not to enforce a classical enterovirus canyon pose, but to let the model insert the ligand on the accessible surface of uncleaved VP0 and at interprotomer junctions, where early-stage capsid binders often sit in related systems (16). It was previously shown that POS inhibits PeV-A3 entry by likely binding the viral capsid (14) therefore the first step was to determine the preferred POS binding site on US WI-09 and 9C-CMKC isolates. Boltz-2 returns a predicted complex together with confidence scores (18). Like AlphaFold3, Boltz-2 includes a confidence module that scores each predicted complex. We used the protein-ligand interface predicted TM (ligand_ipTM in Table 1), which ranges from 0 to 1 and reflects the model’s confidence in the predicted position of POS on the capsid protein surface. Higher values indicate a more reliable placement, and we used this score to rank the predicted POS binding poses.

**Table 1.** Boltz-2 confidence metrics for the predicted POS capsid complexes. complex_plddt (per-residue placement confidence), ligand_ipTM (confidence in POS placement at the protein-ligand interface), confidence_score (overall model confidence), ipTM (protein-protein interface confidence), and the predicted POS site. All metrics range 0-1, with higher values indicating greater confidence

| **System** | **complex_plddt** | **ligand_ipTM** | **confidence_score** | **ipTM** | **Predicted site** |
|---|---|---|---|---|---|
| USWI-09 | 0.803 | 0.411 | 0.801 | 0.793 | VP0-VP1 |
| 9C-CMKC | 0.807 | 0.383 | 0.802 | 0.783 | VP3-VP1 |
| US WI-09 POSA P14 | 0.806 | 0.504 | 0.801 | 0.781 | VP3-VP1 |

A representative two-protomer complex was built for each isolate, and their confidence metrics were compared (Table 1). Confidence in the folded particle was effectively identical in both: complex_plddt, which reports how reliably each residue is placed, was 0.803 for US WI-09 and 0.807 for 9C-CMKC, and the overall confidence_score was 0.801 and 0.802 (0.806 and 0.801 for the resistant isolate US WI-09 POSA P14). The protein-protein interfaces were likewise well ordered, with ipTM 0.793 for US WI-09 against 0.783 for 9C-CMKC. Only ligand_ipTM, which reports how confidently POS itself is registered against the capsid surface, separated them: 0.411 for US WI-09 against 0.383 for 9C-CMKC. Because the two isolates differ by at most 0.004 in complex_plddt, 0.001 in confidence_score, and 0.010 in ipTM<u>, *the models are not distinguishable in how confidently the capsid itself is built; what differs is*</u> *<u>where POS is placed on it, the VP0/VP1 face in US WI-09 and a VP3/VP1 site in 9C-CMKC (</u>Table 1; Figures 2 and 3)*. In the US WI-09 complex, the two POS copies did not collapse into one buried hole (Figure 2). One protomer contacted a VP0 patch spanning residues 46–78, including D46, D47, S49, L51, G53, P54, N55, S56, T73, A71, and T78. The second protomer contacted the same VP0 patch and bridged it to the neighboring VP1 edge, reaching the aromatics Y49 and F50 together with K46, N48, and L51. Shared VP0 contacts across ligand copies (D46, S49, L51, G53, P54, S56, T73, T78) mark a polar hotspot on uncleaved VP0. The laboratory resistance position VP0 K66 sits in this same N-terminal span (14). Even when K66 is not direct contact in every pose, it is close enough that K66R can change local electrostatics and pocket shape. That is the first structural link between the predicted USWI-09 face and experimental escape genetics.

**Figure 1.**
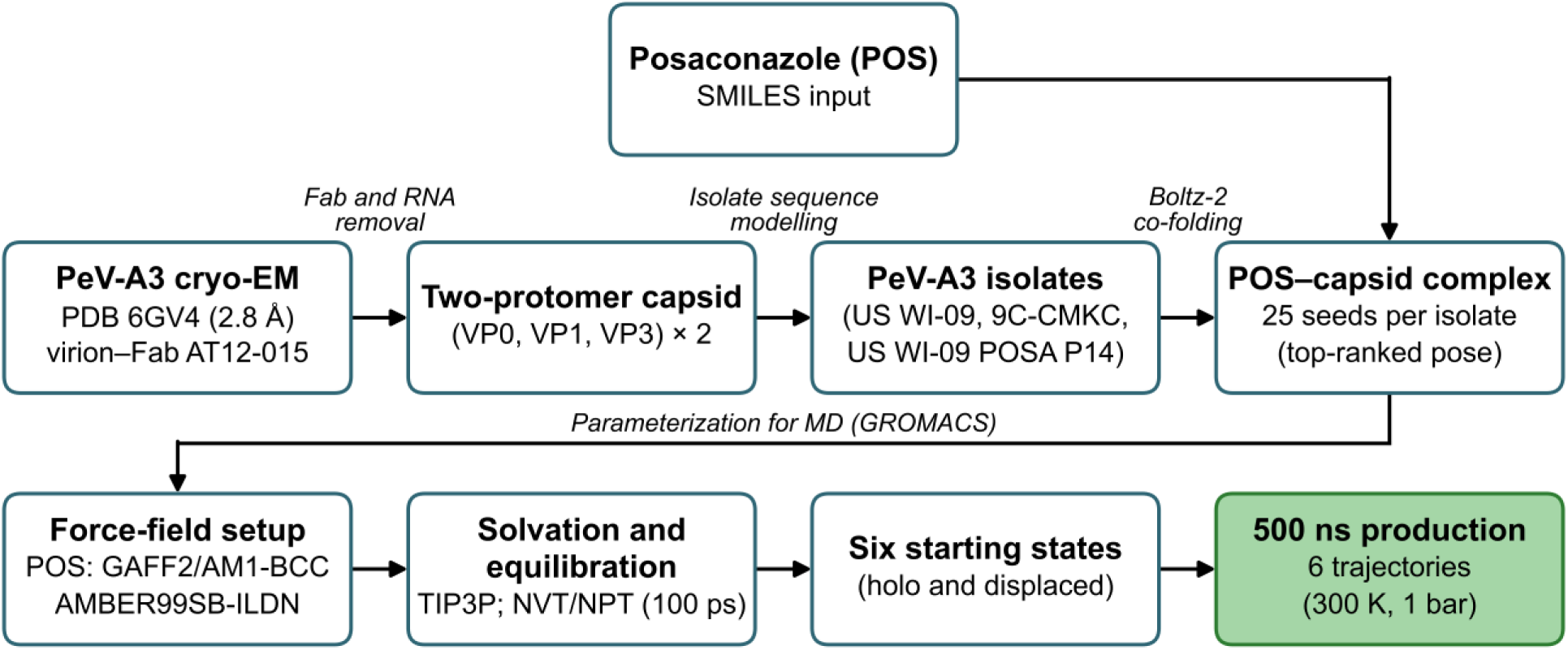
Workflow for constructing and simulating posaconazole-bound PeV-A3 capsid systems.

**Figure 2.**
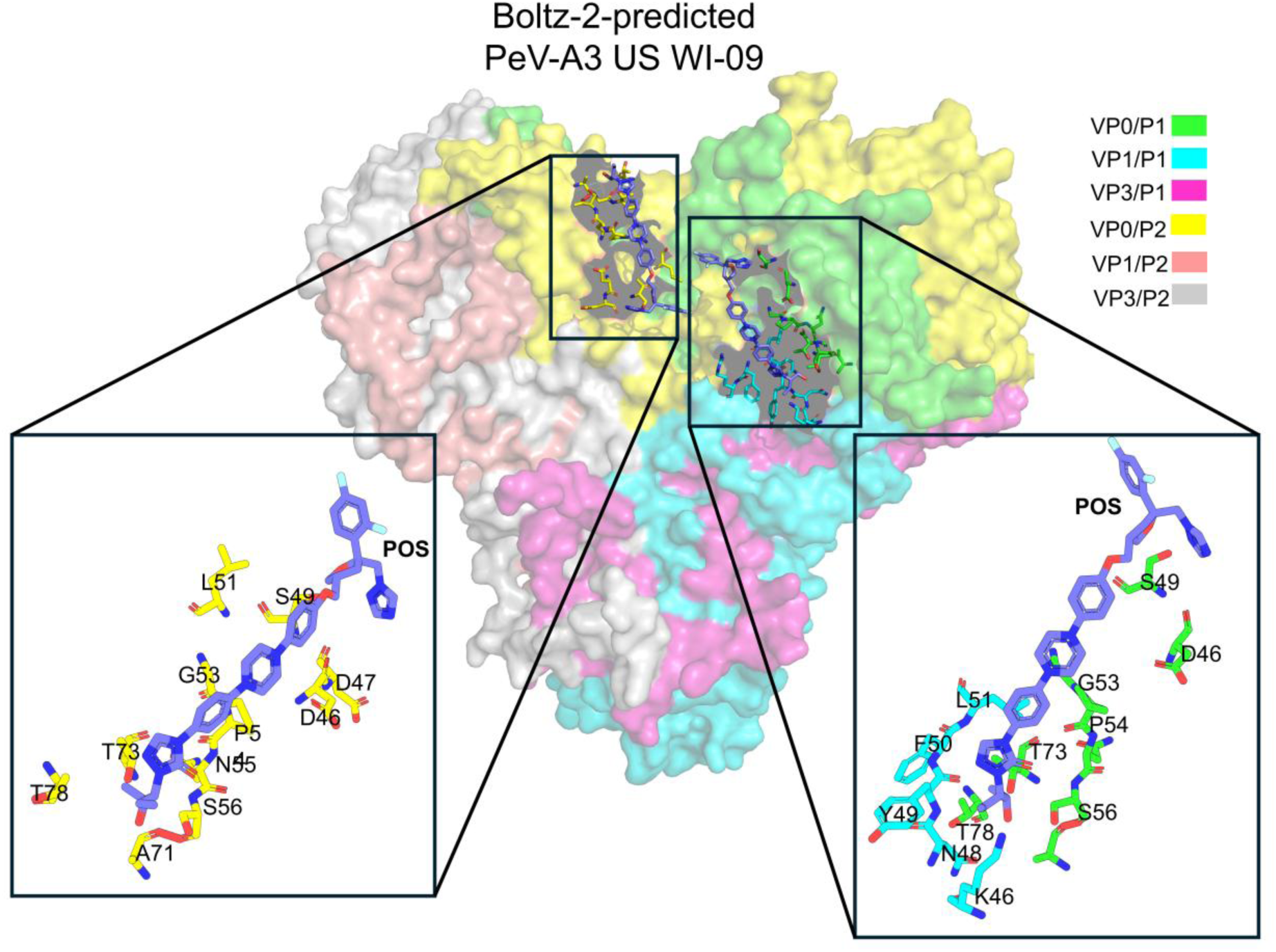
Predicted two-protomer complex of PeV-A3 US WI-09 with POS. Surface view of the two-protomer unit (chains colored as in the legend) with the two POS copies boxed and shown as sticks in the insets. One copy binds the un-cleaved VP0 surface (left: VP0 residues D46-T78), and the second bridges VP0 to the neighboring VP1 edge at the aromatics Y49 and F50 (right), defining the VP0-VP1 interprotomer face.

**Figure 3.**
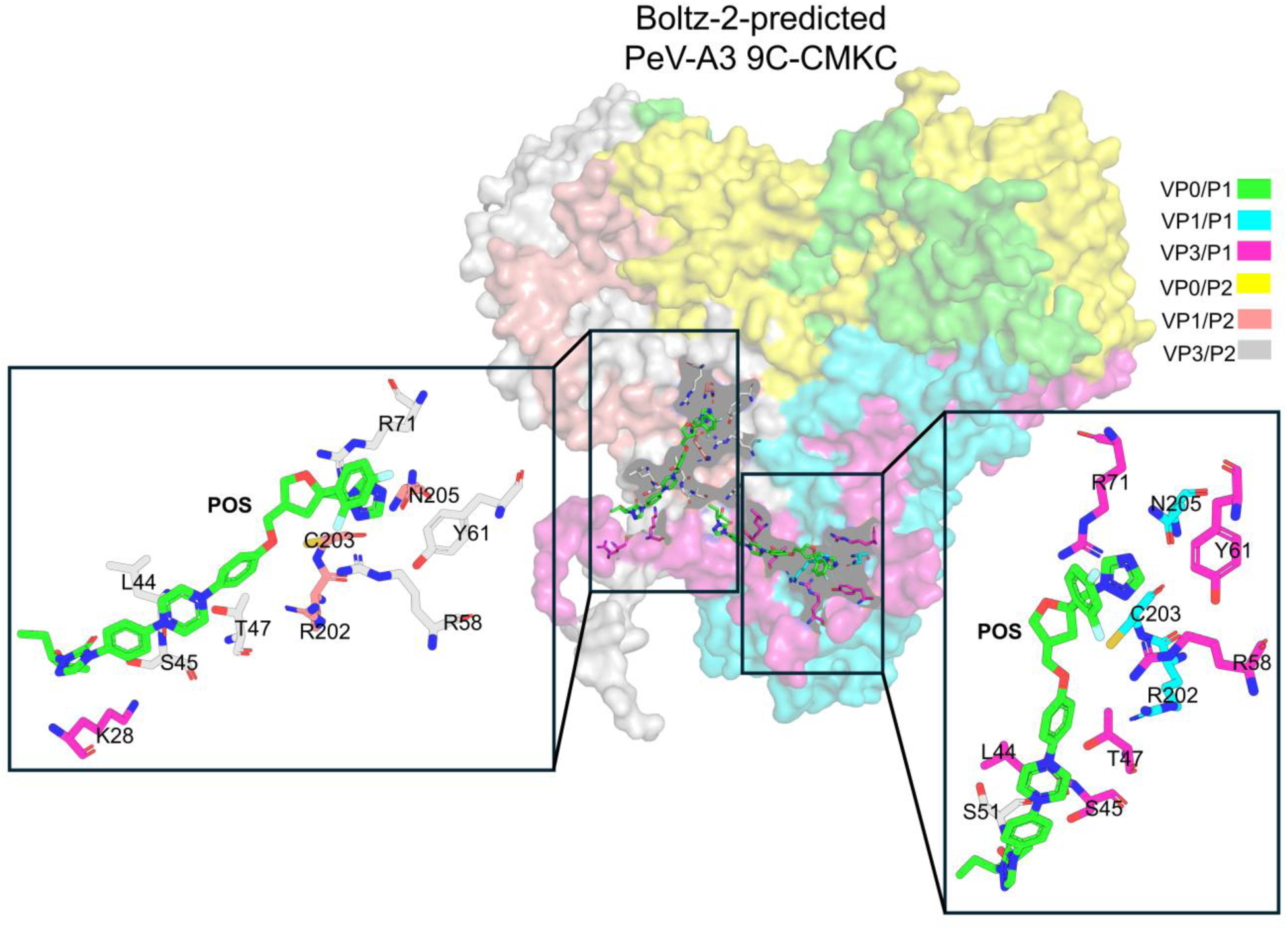
Predicted two-protomer complex of PeV-A3 9C-CMKC with POS. Both POS copies occupy a VP3-centered groove (L44, S45, T47, R58, Y61, R71) with VP1 rim contacts (R202, C203, N205) and no VP0 contacts, placing the ligand at a VP3-VP1 site rather than the VP0-VP1 face seen in US WI-09.

Thereafter, the same two-protomer Boltz-2 setup was run on isolate 9C-CMKC and compared with the four-mutation POS resistant isolate US WI-09 POSA P14. Both keep high complex_plddt and confidence_score, but POS binding is no longer with the VP0-VP1 face. On the US WI-09 POSA P14, POS occupies the VP3 N-terminal residues A18, S19, Y22, K23, and W24 together with T47 (SI Figure 1). VP1 contacts move to R202/C203 rather than VP1 Y49/F50. Ligand_ipTM is 0.504, with confidence_score 0.801 and complex_plddt 0.806 (Table 1). The fold around the new site is still locally confident; the pocket geography is what changed. Isolate PeV-A3 9C-CMKC reaches the same geographic conclusion from natural sequence rather than engineered escape mutations (Figure 3). In the representative two-protomer complex, both POS copies sit in a VP3-centered groove with VP1 rim contacts and no labeled VP0 contacts. One protomer packs against a VP3 annulus that includes L44, S45, T47, R58, Y61, and R71, with VP1 contacts at R202, C203, and N205, and reaches across the interface to K28 of the second protomer. The second protomer uses the same VP3 rim on the opposing protomer (L44, S45, T47, S51, R58, Y61, R71) and the same VP1 edge at R202, C203, and N205. Shared anchors with the resistant model are clear: VP3 T47 recurs, and VP1 again engages the R202/C203 instead of the US WI-09 Y49/F50 aromatics on the VP0-VP1 face. As noted above, the particle fold and protein-protein interfaces of the 9C-CMKC model are as well ordered as those of US WI-09 (Table 1). Ligand_ipTM is the lowest in the POS series (0.383), which reports higher uncertainty in how the flexible ligand registers against that relocated face, not rejection of the complex. Relative to US WI-09, isolate 9C-CMKC therefore abandons the uncleaved VP0. The Boltz residue map for POS therefore has two layers. A <u>functional VP0 core</u> (S49, L51, N55, S56, T73), and a <u>secondary VP3 annulus set</u> (L44, T47, R58, Y61, joined by W24 and the N-terminal arm in US WI-09 POSA P14) appears when that face is mutated or altered by isolate sequence.

### 3.2. MD reveals both stable and transient residue-level interactions with the drug

Predicted bound poses of POS with PeV-A3 capsid were tested in 500 ns all-atom MD of the two-protomer assembly (six chains) with one POS molecule. Each isolate was run as: (1) holo configuration, starting with POS bound to the predicted pocket, and (2) an out-of-pocket (apo-like) configuration, with POS ∼10 Å away from the pocket. This paired simulation helped us discern if the holo configuration is stable and solvent accessible. A pre-docked holo configuration could show artifact trapping of the drug which is un-realistic if there is no true access path to that buried pocket for that drug to travel along. Protein parameters were AMBER99SB-ILDN, ligand parameters GAFF2/AM1-BCC, solvent TIP3P, and the box a rhombic dodecahedron with 1.2 nm padding, after NVT/NPT equilibration (34, 35, 38, 59). The comparison below pairs each residue snapshot with its Rg, RMSD, RMSF, and SASA panel.

Pockets were defined by occupancy rather than by a single frame. A residue enters the pocket if any of its heavy atoms lies within 4 Å of POS in at least 50% of the trajectory (occ50). Around that pocket we track a 12 Å shell, the set of residues within 12 Å of the ligand, as a measure of burial. This section treats the pocket as a whole through two per-frame quantities: engagement, the fraction of pocket residues in contact with POS in a given frame, and the largest connected component (LCC) fraction, the share of pocket residues forming one connected cluster when side-chain centroids within 6 Å are joined, which reports whether the pocket is an intact cavity or a loose collection of side chains.

In all six runs the capsid itself was stable. Capsid RMSD stayed between 0.2 and 0.5 nm, the radius of gyration varied by less than 0.05 nm around means of 3.799 to 3.820 nm, and the solvent-accessible surface area was flat near 550 nm² throughout (Figures 4c to 4f, 5c to 5f; SI Figures 2 to 5). Backbone fluctuation was confined to the N-terminal arms, the inter-barrel loops and the C-termini, with the β-barrel cores below 0.3 nm. The differences between systems are therefore in the ligand, not in the particle.

**Figure 4.**
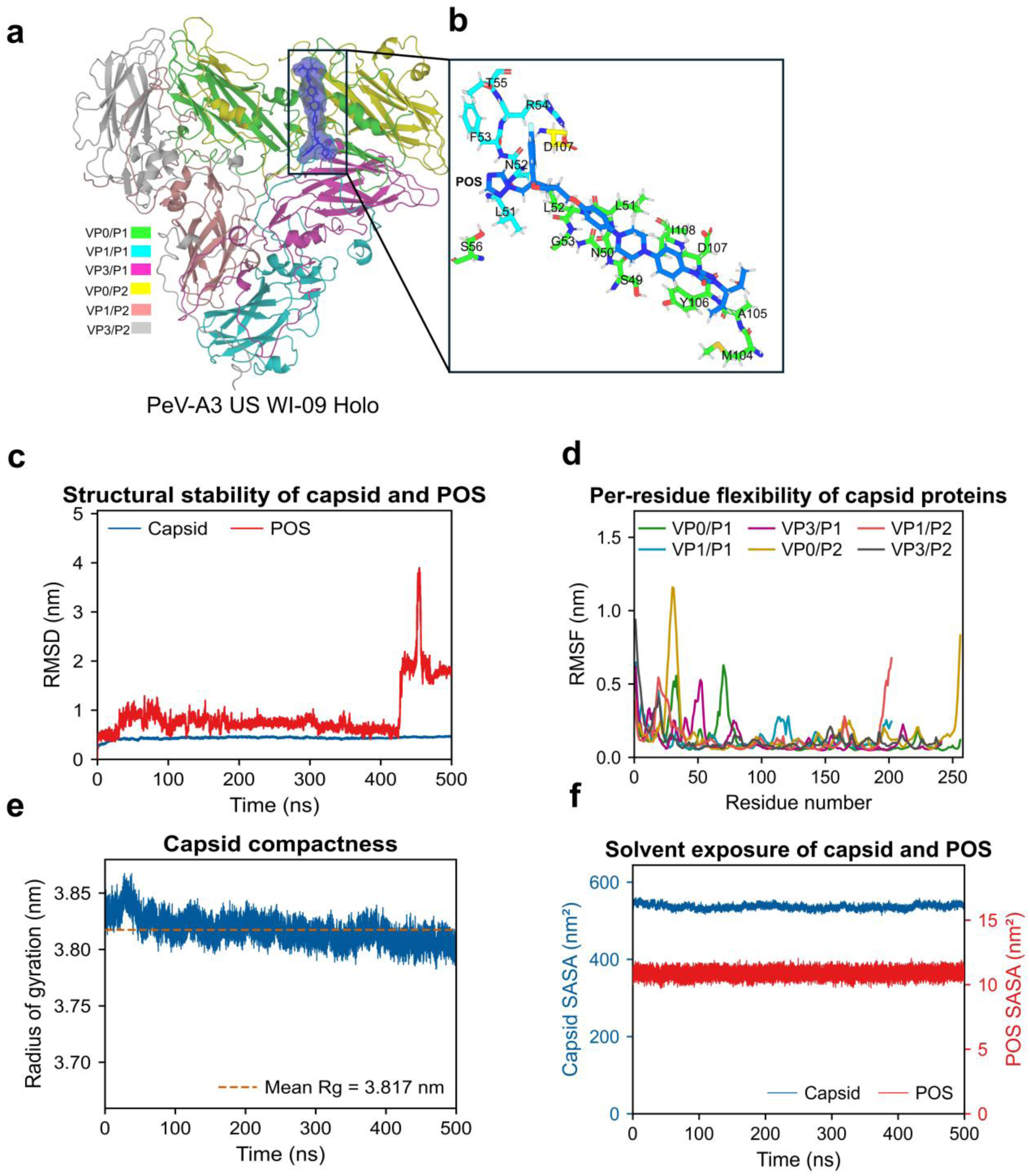
POS binding and stability in PeV-A3 US WI-09 (holo). a) Two-protomer structure at t = 500 ns, chains colored as in the legend, with POS shown as spheres. b) Close-up of the POS pocket at the VP0-VP1 interface; POS packs against the VP0 S49-G53 wall and the VP1 L51-T55 segment within protomer 1, and also contacts VP0 of the second protomer (M104, A105, I106, D107, I108), making this an inter-protomer site. Contacting residues are shown as sticks and labeled. c) Backbone RMSD of the capsid and of POS over 500 ns; the late rise reflects the single dissociation-rebinding event near 450 ns. d) Per-residue Cα RMSF by chain. e) Capsid radius of gyration (mean Rg = 3.817 nm, dashed). f) Solvent-accessible surface area of the capsid and of POS.

#### 3.2.1. POS binds a stable VP0-dominated inter-protomer pocket in US WI-09

In US WI-09 the POS pocket is formed entirely within protomer 1 and spans the VP0-VP1 interface (Figure 4a). It comprises VP0 S49, N50, L51, L52 and G53 together with VP1 Y49, F50, L51, F53, R54 and T55. This is the site predicted by Boltz-2, which place POS at the VP0-VP1 interface. The pose at 500 ns shows the same two walls in direct contact with the ligand, VP0 S49 to G53 with S56 on one side and the VP1 L51 to T55 segment on the other (Figure 4b). To capture the full environment around the bound drug rather than only its direct contacts, a 12 Å shell was defined as all residues with any atom within 12 Å of POS, and each residue was scored by its occupancy, the fraction of the 500 ns trajectory in which it stays inside that shell (1.0 means every frame, 0.5 means half). POS remains in the pocket for the full 500 ns. Its 12 Å shell holds 78 residues occupied in at least half the frames and is dominated by VP0. Part of the outer wall comes from VP0 of protomer 2, through D107 (present in 97% of frames), K113 (92%), T109 (84%), A105 (79%), S116 (65%) and Q271 (55%). POS thus sits within protomer 1 while packing against protomer 2, making this an inter-protomer interface site; in the final frame, D107 and A105 of the second protomer are in direct contact with POS (Figure 4b).

POS is in contact from the first frame and the site is complete within 50 ns, so binding needs no relocation phase. The pocket keeps its shape throughout, breathing between 12.3 and 12.7 Å, so POS does not remodel the site it occupies. Binding stays loose, and the residue-residue network does not tighten as ligand contacts increase. Around 445-459 ns POS briefly leaves the pocket and rebinds: engagement drops below 0.2 in 4.4% of frames, the center-to-center distance reaches 25-32 Å, and the ligand RMSD jumps from ∼0.5 to 3.8 nm before returning partway to 1.5-2.0 nm (Figure 4c, SI Figure 6a). This single dissociation-rebinding event shows that binding here is reversible and loosely held, though one event in one trajectory does not define an off-rate. After displacement of POS by about 10 Å, the ligand returned to the VP0 site. The resulting pocket comprises 19 residues, 16 of them in VP0, and retains all five VP0 residues of the holo pocket (S49-G53) together with VP1 Y49. Compared with holo, it extends deeper into VP0 and engages less of VP1, giving the most VP0-dominated shell of any system (50 of 77 residues from chain A). The final frame shows this directly: the VP0 wall runs from T42, A45 and N50 through L51, L52, G53 and P54 to S56, F57, A58, V69, Q70, A71 and L80, with VP1 contributing only P47, N48 and Y49 (SI Figure 2b). Capture occurred between 90 and 150 ns, with smoothed engagement crossing 0.5 at 87 ns and 0.75 at 154 ns, and POS remained bound thereafter (SI Figure 6b). Recovery of the same site from an independent starting pose identifies VP0 as the preferred US WI-09 site rather than an artifact of the input pose. *<u>Together, the holo and displaced trajectories converge on the same answer: in US WI-09 POS binds</u> <u>a VP0-dominated pocket at the protomer 1-protomer 2 interface, held stably but loosely, and this site is</u> <u>recovered independently of the starting pose</u>*.

#### 3.2.2. 9C-CMKC and the US WI-09 POSA P14 bind the same VP3 escape site, but by opposite mechanisms: an induced-fit lid versus a pre-ordered clamp

Unlike US WI-09, where POS occupies the VP0-VP1 functional core site, both 9C-CMKC and *US WI-09 POSA P14* place POS on the VP3 N-terminal escape region, and in neither system does VP0 contribute to the pocket. However, the two capsids reach that site in very different ways. In 9C-CMKC the pocket is built only from VP1 and VP3 and bridges the two protomers (Figure 5a). It has 16 residues: VP1 D59, I60, F61 and T62 of protomer 1; VP1 T62, R67, D69, N70 and C203 of protomer 2; VP3 V43, L44, R58, A59, F60 and Y61 of protomer 2; and VP3 W24 of protomer 1. No VP0 residue is present, and none of the resistance positions reach the drug or couple to it. The final frame reproduces this arrangement, with the VP1 D59 to T62 loop, R67 and D69, the VP3 V43, R58 to Y61 face and W24 all in contact with POS (Figure 5b). This pocket overlaps the escape site predicted by Boltz-2 for 9C-CMKC and US WI-09 capsids, which comprises VP3 W24, L44, R58 and Y61 with VP1 D69 and C203; all six residues are present in the 9C-CMKC MD pocket. POS in 9C-CMKC therefore sits in the resistant-like site, not the US WI-09 functional core site.

**Figure 5.**
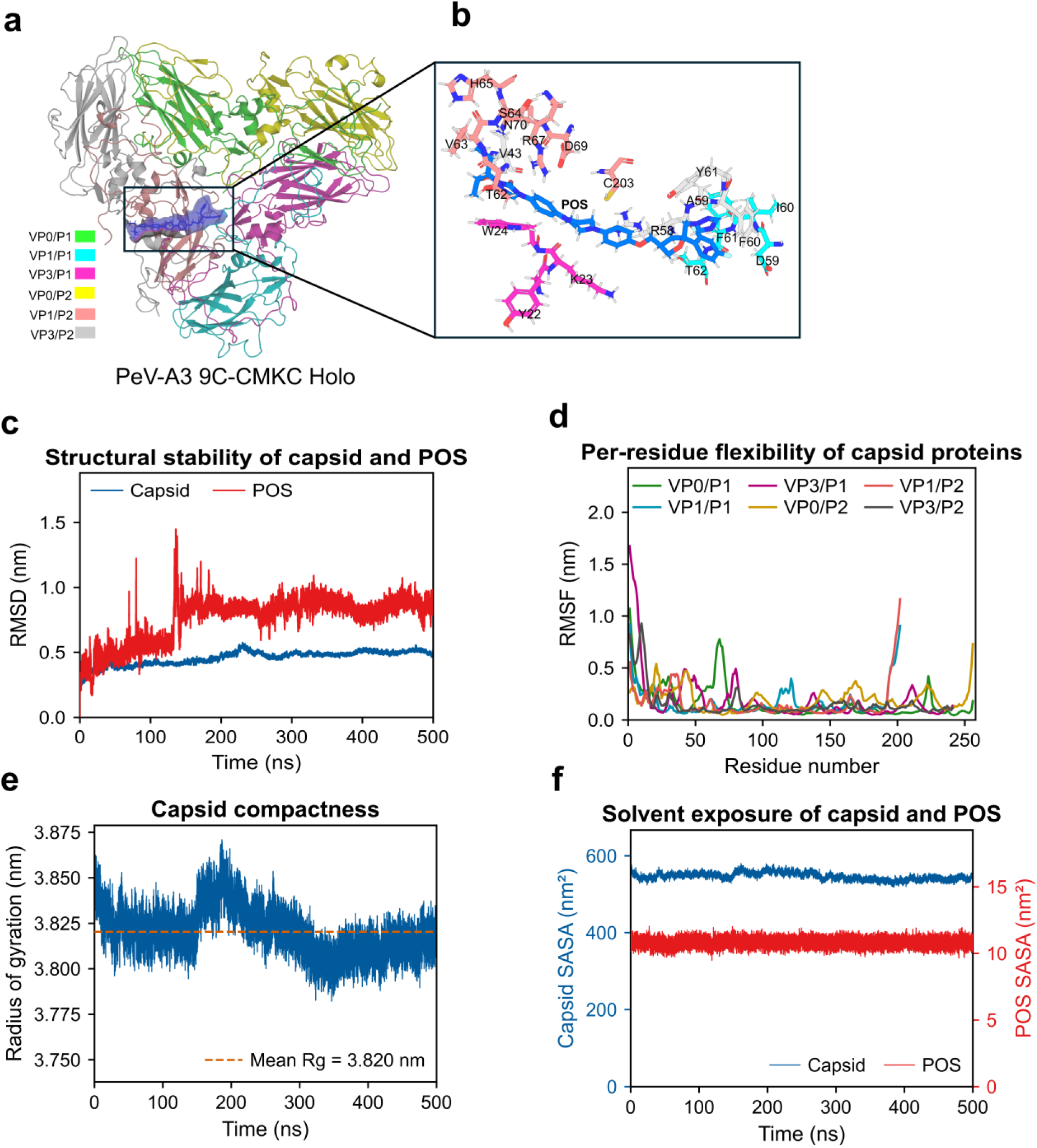
POS binding and stability in PeV-A3 9C-CMKC (holo). a) Two-protomer structure at t = 500 ns, chains colored as in the legend, with POS shown as spheres. b) Close-up of the POS pocket, a VP3-dominated groove with VP1 rim contacts and the VP3 N-terminal arm (W24) closing over the ligand; contacting residues are shown as sticks and labeled. c) Backbone RMSD of the capsid and of POS over 500 ns. d) Per-residue Cα RMSF by chain; the peak at the VP3 N-terminus marks the flexible arm. e) Capsid radius of gyration (mean Rg = 3.820 nm, dashed). f) Solvent-accessible surface area of the capsid and of POS.

Binding proceeds in two phases separated by a transition near 200 ns, where smoothed engagement first exceeds 0.75; beyond 250 ns engagement stays at or above 0.75, a stable bound state rather than a transient encounter (SI Figure 6a). The ligand RMSD follows the same pattern, climbing from about 0.4 nm through a transient excursion near 120 ns to a plateau of about 0.8 to 1.0 nm after 200 ns (Figure 5c). POS inserts by wedging: as it becomes buried, contacts between pocket residues fall rather than rise (LCC 0.30 to 0.22; SI Figure 6c), so POS slots between side chains and replaces residue-residue contacts instead of filling a pre-formed cavity, spanning both protomers and contacting VP1 and VP3 of each. The ligand is closed in by the VP3 N-terminal arm of protomer 1, which becomes steadily more rigid toward its tip (Cα RMSF falling from about 11.9 Å near the N-terminus to roughly 3 Å at W24; Figure 5d). The arm thus acts as a lid that closes once POS is in place, not as part of a pre-formed site.

Displacement by about 10 Å reproduced the 9C-CMKC site, including the VP1 D59 to T62 loop, the VP3 N-terminal arm (Y22, K23, W24) and VP3 L44, with the VP1 loop contributed by the opposite protomer and VP3 W201 added. Neither the pocket nor its shell contains any VP0. The mechanical response is the reverse of holo: POS compacts the site, and contacts between pocket residues rise rather than fall (SI Figure 6d). Over the final 100 ns the ligand migrates within the pocket, releasing the VP3 arm at W24 and S19 and settling onto VP1 I60 and T62 and VP3 W201, with engagement declining from about 0.85 to about 0.35 (SI Figure 6b). The 500 ns pose shows this end state, with I60, T62 and W201 engaged and W24 no longer in contact (SI Figure 3b), and the ligand RMSD steps from 0.5 to 1.5 nm at 70 ns and drifts toward 2.5 nm over the last 100 ns (SI Figure 3c), a sub-pose change rather than dissociation, since engagement never reaches zero.

The US WI-09 POSA P14 binds the same region but holds POS in the opposite way. POS binds the VP3 N-terminal annulus, and VP0 is absent entirely. The pocket has 13 residues: the VP3 N-terminal arm of protomer 1 (L16, T17, A18, S19, Y22, K23, W24), VP3 V43 and L44 of protomer 1, VP1 V63, R67 and N70 of protomer 1, and VP3 V55 of protomer 2. Its 12 Å shell is drawn almost entirely from VP3 and VP1 of both protomers, with no VP0 in the ligand environment. This is the predicted escape site, the VP3 N-terminal annulus with W24 as the universal escape residue, so in the US WI-09 POSA P14, POS again sits where resistance places it, at a site that contains no VP0. POS stays bound for the full 500 ns, and the pocket network stays intact around it. The LCC is the highest of the three systems and rises over the run (SI Figure 6c), and internal packing holds up regardless of how tightly POS engages. This is the opposite of 9C-CMKC, where tighter engagement pulls the pocket residues apart: POS packs against a pre-formed cluster in US WI-09 POSA P14 but wedges that cluster open in 9C-CMKC. POS is also the least buried of the three ligands and sits off-center, anchored at the surface of the cluster rather than enclosed by it. Two features hold it there: an aromatic clamp of VP3 Y22 and W24 that contacts POS in nearly every frame, and VP3 V55 of protomer 2, which contacts POS almost constantly and stays within 6 Å of VP1 N70, VP1 V63 and VP3 V43 of protomer 1, knotting the two protomers together as the pocket hub. Cross-correlation of residue motion with the POS centroid confirms the asymmetry: only the VP3-arm residues (A18, S19, Y22, K23, W24) track the ligand above the noise floor, while VP1 residues move independently of it. Over the last 100 ns the arm tip rearranges while the clamp holds, with S19 taking over as L16 and T17 release; the 500 ns pose shows A18, S19, Y22, K23 and W24 engaged and L16/T17 gone (SI Figure 4b).

Displacing POS by about 10 Å reproduced the US WI-09 POSA P14 site from 10 VP3 residues, again with no VP0. The aromatic clamp re-formed on the opposite protomer (Y22 and W24 from VP3 of protomer 2; SI Figure 5b), identifying the site as the two-fold VP3-VP3 arm cleft rather than a single-arm feature. Capture was immediate and POS did not dissociate, with the ligand RMSD flat near 2.2 nm throughout (SI Figures 5c, 6b). The displaced site is looser than holo, with the LCC and internal packing dropping sharply (SI Figure 6d); POS compensates by moving with the arm rather than being enclosed by it. Isolates 9C-CMKC and US WI-09 POSA P14 therefore reach the same region of the capsid, sharing VP3 W24, V43 and L44 with VP1 R67 and N70 at equivalent positions (on different protomers), and neither pocket contains VP0. What differs is how the arm holds the drug. In US WI-09 POSA P14, the VP3 arm is already ordered when POS binds (Cα RMSF ∼0.9-1.1 Å at Y22-W24), and POS packs against it. In 9C-CMKC the arm is disordered (∼5.0 Å at Y22) and closes over a ligand that has wedged its way in. *<u>The same</u> <u>escape site is thus reached by two opposite mechanisms: a pre-formed clamp in US WI-09 POSA P14, and</u> <u>an induced-fit lid in 9C-CMKC</u>*.

### 3.3. Pocket chemistry analyses reveal POS interaction is compact and reversible in PeV-A3 US WI-09 compared with the redundant, interface-spanning site of isolate PeV-A3 9C-CMKC

Two capsids hold one drug in two different ways. In US WI-09 the pocket is built once and then used; in 9C-CMKC it assembles around the ligand as the trajectory runs. Three measurements resolve this contrast. Contact occupancy is the fraction of the 5,001 frames in which a residue has any heavy atom within 4 Å of POS, and it ranks residues by how reliably they touch the drug (Figures 6a, 7a). The internal contact matrix applies the same measure to pairs of pocket residues, using side-chain centroids within 6 Å, and reports how the pocket is wired to itself rather than to the ligand (Figures 6b, 7b). The timeline resolves contact frame by frame, separating residues present from the start from those recruited later (Figures 6c, 7c). The two capsids reach similar overall engagement by routes that have almost nothing in common. In US WI-09, 7.5 of 11 pocket residues are in contact in an average frame; in 9C-CMKC, 10.5 of 16. The resemblance ends there. *<u>US WI-09 holds POS with a small number of uniformly reliable contacts confined to one</u> <u>protomer and loses the ligand once. 9C-CMKC holds it with a larger number of individually weaker</u> <u>contacts spread across two protomers and never loses it</u>*: not one frame of the 9C-CMKC trajectory is free of pocket contact, against 87 such frames in US WI-09.

**Figure 6.**
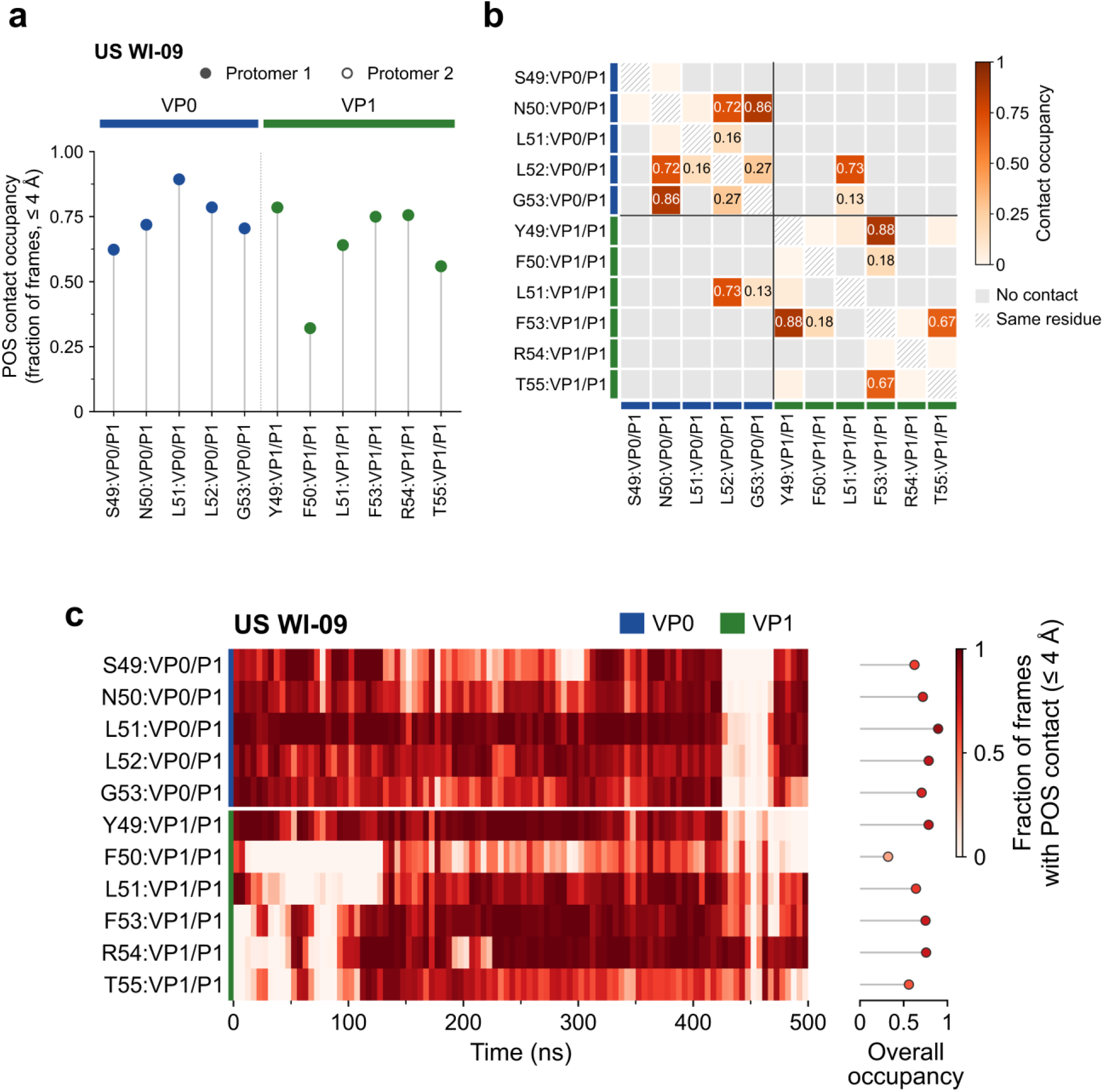
POS contact chemistry of the US WI-09 holo pocket over 500 ns. a) Contact occupancy of each occ50 pocket residue, defined as the fraction of 5,001 frames in which any heavy atom of the residue lies within 4 Å of POS. Residues are ordered by sequence and grouped by capsid protein (VP0, blue; VP1, green); filled symbols denote protomer 1 and open symbols protomer 2. All US WI-09 pocket residues belong to protomer 1. b) Internal contact matrix of the pocket, giving the occupancy of each residue pair whose side-chain centroids lie within 6 Å. Values are printed for occupancy ≥ 0.10; grey indicates no contact and hatching marks the diagonal. Solid lines separate capsid proteins. c) Per-residue contact timeline binned at 5 ns, with color giving the fraction of frames in each bin in which the residue contacts POS. The stem plot at right gives overall occupancy across the full trajectory

Occupancy distributions separate a compact site from a redundant one. The eleven US WI-09 residues form a narrow band, with nine of them between 0.56 and 0.89 (Figure 6a). VP0 L51 is the most persistent contact at 0.89 and makes the closest mean approach in US WI-09 (3.74 Å), with VP0 L52 and VP1 Y49 next at 0.79. Only VP1 F50 is weak, at 0.32, and it is the one US WI-09 residue whose mean minimum distance exceeds 5 Å (5.15 Å), so it lines the rim rather than the site. VP0 averages 0.75 and VP1 0.63, the balance expected of a pocket that straddles an interface instead of belonging to one subunit. The sixteen 9C-CMKC residues spread much more widely, from 0.42 to 0.91 (Figure 7a), and the spread sorts the site into a VP3 core and a VP1 periphery. VP3 Y61 (0.91), VP3 R58 (0.89) and VP1 C203 (0.78) lead the ranking and also give the three closest mean approaches (3.45, 3.54 and 3.80 Å), while the lowest VP1 contacts, D59 and I60 of protomer 1, trail at about 0.5 with mean distances of 5.5 to 5.7 Å. The weakest contact in each system means something different. US WI-09 F50 was always peripheral. 9C-CMKC I60 is low only because it was absent for the first 200 ns; after the transition it holds 0.70.

**Figure 7.**
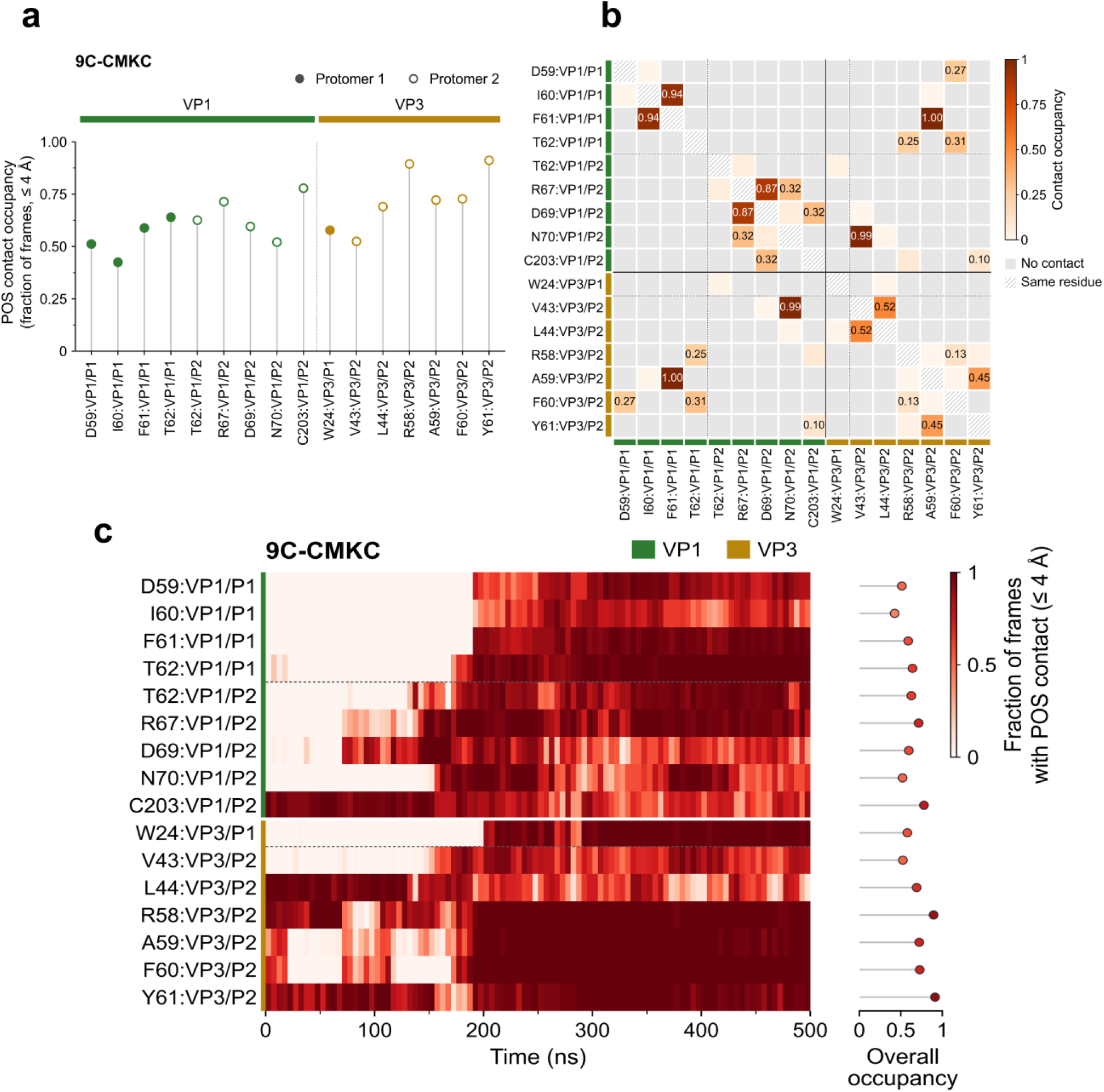
POS contact chemistry of the 9C-CMKC holo pocket over 500 ns. Panels as in Figure 6, for the 9C-CMKC holo trajectory. Pocket residues come from VP1 (green) and VP3 (gold) of both protomers; no VP0 residue meets the occ50 criterion. The timeline in c resolves the transition at approximately 190 to 200 ns, after which VP3 W24, VP3 A59, VP3 F60 and the VP1 D59 to T62 loop of protomer 1 become persistent contacts while the lowest VP1 contacts, D59 and I60 of protomer 1, trail at about 0.5 with mean distances of 5.5 to 5.7 Å. Note in b that the W24 row is almost entirely grey despite the high ligand occupancy of that residue in a and c: W24 contacts POS without contacting any other pocket residue.

The timelines show one pocket completing early and the other assembling in two stages. In US WI-09 the VP0 wall is present from the first frame and the VP1 wall closes within 50 ns (Figure 6c). All five VP0 residues exceed 0.5 smoothed occupancy at t = 0 and hold 0.8 to 0.98 through 400 ns, while VP1 F53 crosses 0.5 at 21 ns, T55 at 24 ns and R54 at 49 ns, climbing from 0.22 to 0.34 in the first 100 ns to 0.85 to 0.98 between 100 and 400 ns. POS begins against VP0 and draws the VP1 half of the site around itself inside the first tenth of the run. 9C-CMKC takes four times as long and recruits in a defined order (Figure 7c). Six residues are engaged at the start (VP1 C203, VP3 L44, Y61, R58 and F60 at t = 0, VP3 A59 at 3 ns); the other ten arrive one at a time over the following 200 ns, as VP1 D69 at 71 ns, VP1 T62 of protomer 2 at 134 ns, VP1 R67 at 140 ns, VP1 N70 at 156 ns, VP3 V43 at 159 ns, VP1 T62 of protomer 1 at 175 ns, VP1 F61 at 190 ns, VP1 D59 at 192 ns, VP3 W24 at 200 ns and VP1 I60 at 215 ns. Comparing the windows before 190 ns and after 200 ns, eight residues gain and two lose. The gains go nearly to completion: W24 from 0.00 to 0.96, VP1 D59 from 0.00 to 0.83, VP1 F61 from 0.00 to 0.95, VP1 T62 of protomer 1 from 0.08 to 0.98, VP3 A59 from 0.27 to 1.00 and VP3 F60 from 0.28 to 1.00. The two losses are the residues that held the ligand at the surface, VP1 C203 from 0.95 to 0.67 and VP3 L44 from 0.92 to 0.55. The initial anchors are demoted, not displaced.

The internal matrices explain why one site releases the drug and the other does not. The US WI-09 pocket is two cliques joined by a single bridge (Figure 6b). Within VP1, Y49 to F53 is occupied in 0.88 of frames and F53 to T55 in 0.67; within VP0, G53 to N50 is 0.86 and L52 to N50 is 0.72. Exactly one pair crosses between the proteins at appreciable occupancy, VP1 L51 to VP0 L52 at 0.73, with VP1 L51 to VP0 G53 a distant second at 0.13. Every pair lies within protomer 1, and the pocket contains no inter-protomer contact at all. A single hinge holding two otherwise independent walls is a loose architecture, and it matches a ligand that sits 7.4 ± 1.1 Å from the pocket center and leaves once during the run. The 9C-CMKC pocket is stitched across the interface instead (Figure 7b). Its strongest pair, and the strongest in either matrix, crosses the protomer boundary: VP3 A59 of protomer 2 to VP1 F61 of protomer 1, occupied in every frame. VP3 V43 of protomer 2 to VP1 N70 of protomer 2 follows at 0.99, VP1 F61 to I60 at 0.94 and VP1 D69 to R67 at 0.87, and three further cross-protomer pairs appear at lower occupancy, all of them VP3 of protomer 2 reaching VP1 of protomer 1 (F60 to T62 at 0.31, F60 to D59 at 0.27, R58 to T62 at 0.25). *<u>Redundancy of</u> <u>this kind, rather than the strength of any one contact, is what keeps POS in the 9C-CMKC site</u>*.

The dissociation event in US WI-09 is selective, and the returned pose is not the original one. Mean engagement over 427 to 460 ns falls to 0.22, but the loss is carried almost entirely by VP0: S49 drops to 0.00, N50 to 0.01, G53 to 0.04, L51 to 0.08 and L52 to 0.09, while VP1 R54 holds at 0.70 and VP1 L51 at 0.45 (Figure 6c). The ligand peels off the VP0 floor and stays tethered to the VP1 edge by the half of the pocket that the internal matrix shows to be most strongly self-connected. Rebinding after 460 ns restores the VP0 wall (L51 0.78, L52 0.74, N50 0.72) but not VP1 Y49, which falls from 0.88 before the event to 0.05 after it while R54 rises to 0.93. The ligand returns to the same floor against a shifted VP1 contact set, which is why the ligand RMSD settles near 1.5 to 2.0 nm rather than returning to its pre-event value. Nothing of this kind occurs in 9C-CMKC, where every residue that is gained at the transition is still in place at 500 ns.

VP3 W24 is a lid, not a wall. The single clearest result in these panels comes from reading the occupancy and the matrix together for one residue. W24 contacts POS in 0.96 of frames after the transition and in every frame of the last 100 ns, yet its highest internal contact with any other pocket residue is 0.017, with VP1 T62 of protomer 2, and most of its row is exactly zero (Figure 7b). W24 is connected to the ligand and to nothing else in the site. It arrives last, lands on the drug and makes no lateral contacts once it is there, which is what a closing lid looks like in this representation and what distinguishes it from every US WI-09 pocket residue, each of which is embedded in a local contact clique. This is the structural form of the escape residue identified by the resistance analysis. *<u>That the resistant-type 9C-CMKC site holds POS more</u> <u>redundantly than the reversible US WI-09 pocket shows that resistance here relocates the drug to a tighter</u> <u>but functionally irrelevant site, rather than weakening how it binds</u>*.

### 3.4. Resistance positions: local chemistry and coupling to the drug site

From previous literature, we know that four positions confer POS resistance in US WI-09 POSA P14: VP0 K66R, VP0 A124T, VP3 N88S and VP1 Y224C. US WI-09 carries the parental residue at all four, and 9C-CMKC has a T124 and R224 naturally while retaining K66 and N88. Because the three capsids bind POS at different sites, the same four positions sit in different relationships to the drug in each. Of the four, only VP0 K66 has any structural relationship to the bound drug; the rest of this section is the evidence for that claim and for what the other three do instead. Only one position comes near the ligand, and only in US WI-09. K66 occupies the 12 Å POS shell, the residues lying within 12 Å of the drug, in 18% of frames, while the other three stay 27 to 61 Å away and never enter it (Table 2, Figure 8a). Once engaged, its distance to POS tracks pocket engagement, measured by the Pearson correlation coefficient r, which runs from −1 to +1 and is negative here because K66 moves closer as POS binds more fully: r shifts from +0.32 in the first 100 ns to −0.68 in the last block (−0.39 overall). The displaced run makes the same point without the input pose to explain it, with K66 in the shell for 99.6% of the final 150 ns and r = −0.50 (Figure 8b). K66 contacts POS directly in only 0.14% of frames, so it sits in the second shell behind the V69 to A71 wall. In 9C-CMKC, K66 stays 38 Å or more away, and the position that approaches instead is VP1 R224 of protomer 2, which reaches the ligand shell late in the run. Each of these two capsids therefore has one resistance position near the drug, and in both cases it belongs to the protein that forms that capsid’s pocket: K66 in VP0 for the VP0-VP1 site, R224 in VP1 for the VP1-VP3 site. None approaches POS in the US WI-09 POSA P14.

**Figure 8.**
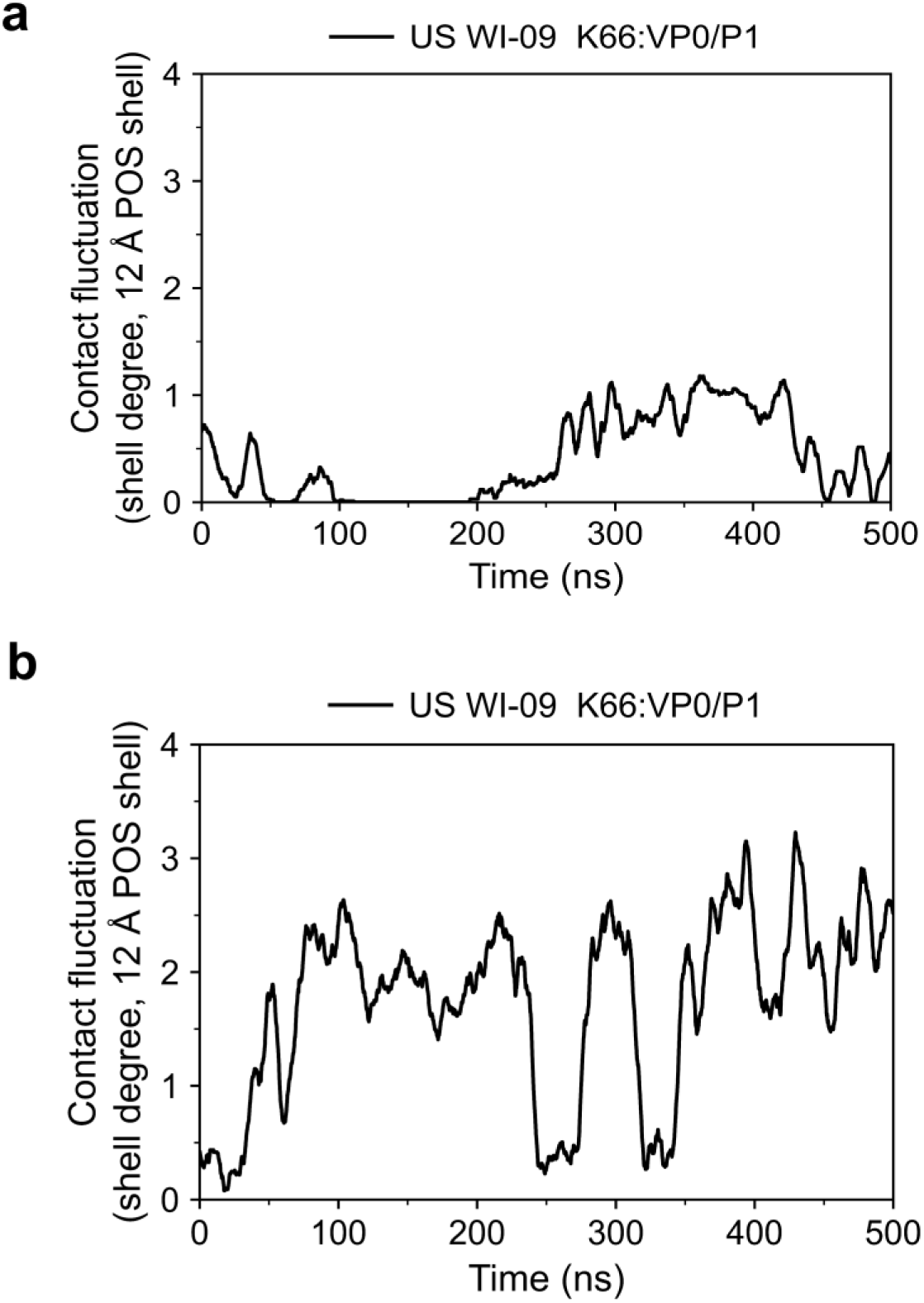
Shell-degree fluctuation of VP0 K66 over 500 ns in PeV-A3 US WI-09. Shell degree is the number of contacts K66 makes within the 12 Å POS shell, smoothed with a centered 10 ns rolling mean. a) Holo run: K66 is absent from the ligand shell for the first 200 ns, after which it engages intermittently. b) Displaced run: K66 enters the shell within the first 50 ns and remains engaged for most of the trajectory, reaching higher degree than in holo. The remaining three resistance positions never enter the shell in either run or are not shown.

**Table 2.** VP0 K66 proximity and POS pocket engagement by 100 ns interval in US WI-09 holo*ᵃ*.

| **Interval (ns)** | **Pearson *r*** | **Mean K66-POS distance (Å)** | **Mean POS engagement** | **K66 in 12 Å POS shell (% frames)** |
|---|---|---|---|---|
| 0-100 | +0.32 | 17.7 | 0.57 | 0 |
| 100-200 | +0.21 | 19.5 | 0.73 | 0 |
| 200-300 | −0.05 | 14.5 | 0.77 | 8 |
| 300-400 | −0.35 | 11.8 | 0.83 | 59 |
| 400-500 | −0.68 | 15.9 | 0.52 | 21 |
| Full trajectory | −0.39 | 15.9 | 0.69 | 18 |

<sup>*a*</sup> US WI-09 holo trajectory (500 ns, 5,001 frames); K66 refers to VP0 of protomer 1, capsid numbering. <sup>*b*</sup> Pearson correlation between the K66-POS distance and POS engagement; negative values indicate that K66 lies closer when engagement is higher. The full-trajectory value is computed over all frames, not averaged across intervals. <sup>*c*</sup> Minimum heavy-atom distance to POS. <sup>*d*</sup> Fraction of pocket residues within 4 Å of POS. <sup>*e*</sup> Percentage of frames with K66 within 12 Å of POS.

None of the other three touches the drug, so their effects must be chemical or allosteric. Of these, position 224 changes most. In US WI-09 protomer 1, Y224 is anchored on VP3 by four hydroxyl hydrogen bonds, to Q124 (0.58 of frames), S86 (0.46), K99 (0.44) and N118 (0.26), and is rigid (Cα RMSF 1.5 Å). C224 in US WI-09 POSA P14 forms no polar contacts at all; its local degree, the number of residues it contacts, falls from 4.73 to 1.75, and the VP1 tail comes loose (RMSF 4.2 Å; tail profile 1.8 to 6.3 Å across residues 217 to 226, against 1.4 to 1.9 Å in US WI-09). 9C-CMKC R224 also loses every VP3 partner, salt-bridging only within VP1 (E226 at 0.29 and 0.37, E98 at 0.53), with RMSF 4.4 and 5.4 Å. A cysteine and an arginine share no chemistry, yet both abolish the Tyr224–VP3 anchor, which identifies loss of that anchor as the operative change and fits Y224C conferring the strongest resistance (34-fold).

Position 66 moves the opposite way. In US WI-09 protomer 1, K66 bridges VP0 to VP3 intermittently through D187 (0.42) and S189 (0.39) while in protomer 2 it makes almost no salt bridges (0.03) and is mobile (RMSF 4.9 Å). The US WI-09 POSA P14 arginine reverses this exactly where the lysine was loosest: in protomer 2, R66 holds VP3 D187 in 0.93 of frames and VP1 D38 in 0.77 and is rigid at 1.3 Å, installing a stable two-partner electrostatic lock across VP0, VP3 and VP1. In 9C-CMKC the lysine pairs with VP0 E64 (0.23 and 0.55) rather than reaching VP3. The other two positions change little. A124 makes no polar contacts in US WI-09; both resistant capsids carry threonine and gain the same hydrogen bonds to S125 (0.94 and 0.54 in 9C-CMKC; 0.74 and 0.67 in US WI-09 POSA P14) and to the I136 backbone, with no change in mobility, so 9C-CMKC holds one resistance allele before any selection. N88 is the least perturbed: both resistant capsids reproduce its hydrogen-bond network and its inter-protomer contact with VP0 R132 of the opposite protomer (1.00, 0.97 and 0.99). That is itself a result, since N88S confers 12-fold resistance while leaving its environment intact, so it must act through something other than local remodeling, most plausibly that R132 contact. Averaged over both protomers, only 224 loses an interface (degree 3.53 to 1.99, mobility 2.06 to 3.03 Å) while 66 gains one (salt bridges 0.33 to 1.31, mobility 3.34 to 1.78 Å).

Proximity and local chemistry cannot show whether a position influences the pocket at a distance, so dynamic coupling was quantified as the mean absolute Cα-Cα cross-correlation, |C|, between each resistance position and the pocket residues, averaged over five 100 ns blocks, where 0 is independent motion and 1 is motion as one rigid body. Because cross-correlation is distance-dependent, each value was compared with a distance-matched null distribution (2,000 random non-pocket residue sets). Allosteric routes were traced separately as shortest paths through the residue contact network, with each path length ranked against the same distance-matched null, so that a lower percentile means a path shorter than chance. K66 of protomer 1 in US WI-09 gives |C| = 0.130 against a null mean of 0.082, exceeding all 2,000 draws (p < 0.001), with VP1 Y49 as its strongest partner (0.19); it is the only site-pocket coupling in US WI-09 that beats its distance-matched null. In the displaced run it couples twice as strongly as distance predicts (0.244 against 0.120, p < 0.001) and reaches POS in four steps through VP0 D67, V68 and A58. Network paths agree without quite reaching significance: N88 has the shortest route of the four (6.2nd percentile), through VP3 V84, Q236, L144, L192 and T193 into VP1 Y49, and Y224 uses the same relay, so the two positions that share VP3 S86 also share their route into the pocket. All eight paths enter POS through VP1 Y49 or R54. Neither resistant capsid shows any such coupling: in US WI-09 POSA P14, the eight sites connect only by routes of 8 to 19 residues, none shorter than distance-matched controls, and 9C-CMKC shows none at all. The pocket-to-ligand coupling runs the other way: no US WI-09 pocket residue moves with POS above noise, against 8 of 16 in 9C-CMKC, so the capsid with a coupled resistance position is the one whose pocket does not hold the drug as a unit.

One position behaves unlike the other three. *<u>VP0 K66 is the only residue with a direct structural</u> <u>link to bound POS, and only in US WI-09</u>*: it enters the ligand shell, it is the single position whose motion tracks the pocket beyond what distance explains, and when POS is displaced and left to find its own site it returns to VP0 with K66 in its second shell for 99.6% of the final 150 ns. A position defined by resistance selection thus maps onto the site the ligand selects unaided, which identifies the VP0-VP1 interface as the drug-relevant site rather than an artefact of the predicted pose. The other three never approach that site. Position 224 instead releases the VP1 C-terminal tail from VP3, an effect produced equally by US WI-09 POSA P14 cysteine and by the arginine that 9C-CMKC carries naturally, and 9C-CMKC is the capsid in which POS no longer binds the VP0-VP1 face at all. *<u>Resistance can therefore arise two ways: by perturbing</u> <u>the functional site directly, as at position 66, or by relocating the ligand away from it, as at position 224</u>*.

### 3.5. POS treatment reduced viral titer of US WI-09 and 9C-CMKC isolate in Vero-p cells but is unable to reduce the titer of 9C-CMKC isolate in Vero-MYADM cells

From our previous results, we determined that POS has different binding profiles with PeV-A3 US WI 09 binding in a pocket localized to the VP0-VP1 region of the capsid versus VP3-VP1 in the 9C-CMKC isolate. Now, we wanted to evaluate whether this differential binding of POS to viral capsid proteins impact the internalization and subsequent replication of the isolates in the presence of a POS doze (0.3uM) that has been tested previously in Vero-p^14^. We were also interested in evaluating the effectivity of POS at the same concentration in a more permissive cell line Vero-MYADM. Our results demonstrate that POS reduced the mean relative viral titer of USWI-09 and 9C-CMKC by ∼1 log folds and 1.5 log folds respectively in Vero-p cells, 5 days post infection as compared to non-POS treated cells (Figure 9a). In Vero-MYADM cells, POS reduced only the mean relative viral titer for the US WI-09 isolate by ∼2 log fold. For the 9C-CMKC isolate, there was no reduction in viral titer by the end of 5 days post infection in the presence of POS (Figure 9b).

**Figure 9.**
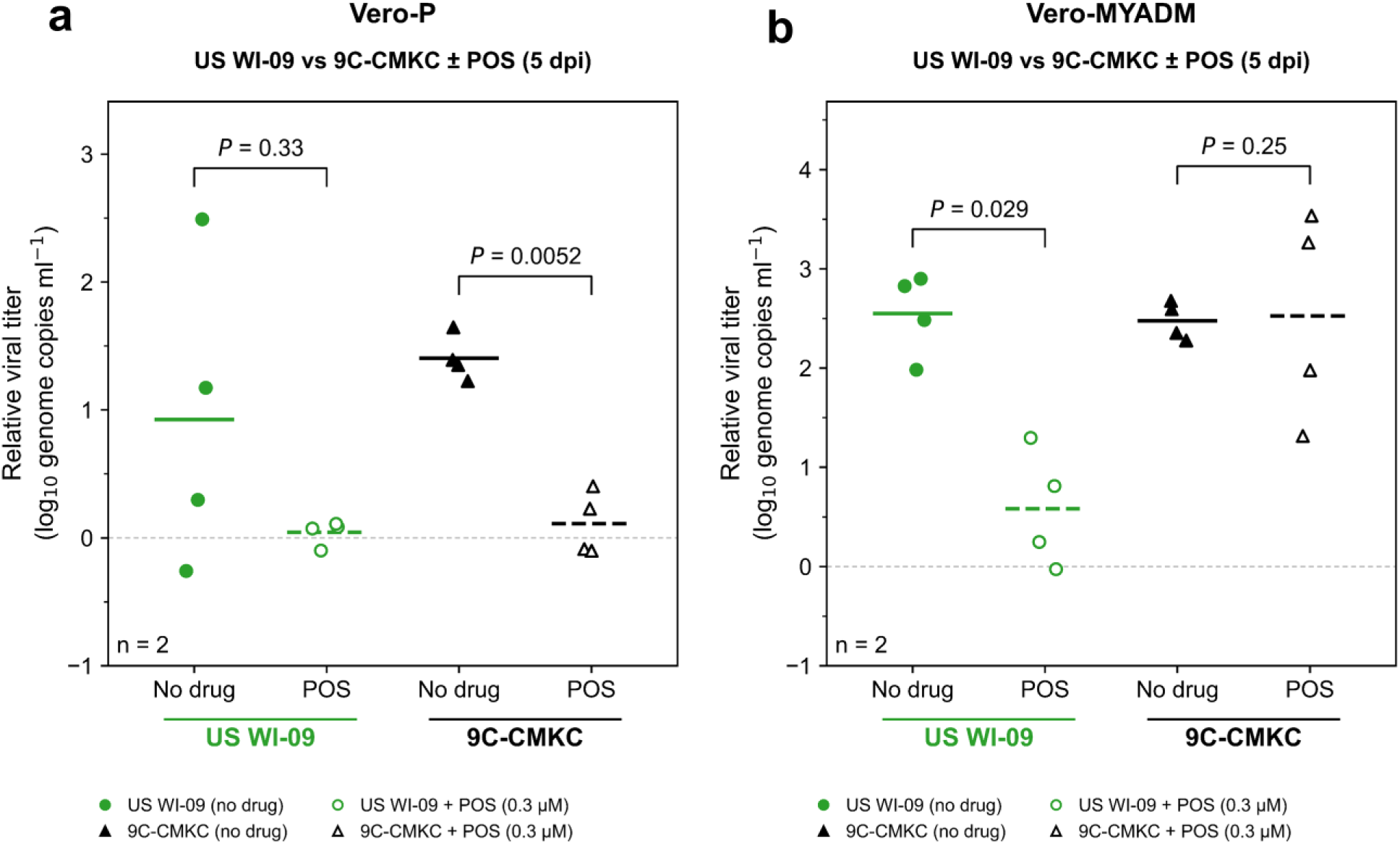
POS treatment reduced viral titer of US WI-09 and 9C-CMKC isolate in Vero-p cells but is unable to reduce the titer of 9C-CMKC isolate in Vero-MYADM cells. Relative viral genome titers (genome copies/ml) measured 5dpi in a) Vero-p and b) Vero-MYADM cells infected with US WI-09 and 9C-CMKC. Cells were treated with POS (0.3 μM) or vehicle control (no drug). Individual data points represent biological replicates, and horizontal lines indicate group means. Corresponding p values between groups reported using two tailed student t-test.

## DISCUSSION

Posaconazole (POS) blocks an early step in the PeV-A3 life cycle by binding onto the viral capsid, yet its capsid target was never localized and its activity against circulating viruses never tested. POS does have a defined binding site in the 2009 isolate US WI-09 - a VP0-dominated pocket at the VP0-VP1 interprotomer face, and 2022 isolate from a different clade does not use that site at all. We built the two-protomer particle with equal confidence for all three sequences (complex_pLDDT ∼0.80; ipTM ∼0.78), so only the ligand site moves: to a VP3 annulus with VP1 R202/C203 contacts in both 9C-CMKC and the laboratory-selected resistant variant (USWI-09 POSA P14). Six unrestrained 500-ns trajectories reproduced that split, and it survives removal of the input pose: displaced ∼10 Å and released, POS recaptured the VP0 site in US WI-09 within 150 ns, while both resistant backgrounds re-converged on the VP3 N-terminal arm. VP0 exclusion is a property of the resistant capsid, not of the starting structure and VP0 is the protein experimentally shown to engage MYADM, the essential PeV-A3 receptor.

Escape is not weakened binding. Binding enthalpy is favorable in all six ensembles, converging to −1.34 to −1.75 kcal/mol per pocket residue with engagement indistinguishable across runs; the 9C-CMKC site is the more redundant of the two, stitching POS across the protomer interface and never releasing it. Our interpretation is that affinity ranking cannot see what matters here. Interaction entropy did not converge in any of the six systems, so a ΔG ordering (9C-CMKC most favorable, US WI-09 formally unbound) inverts the cell data and reflects the instability of that term rather than the chemistry of the complexes. Even with a well-behaved entropy term, though, we would not expect ΔG to identify the productive site, because a POS ligand can bind comparably well at a surface the virus does not need. The per-residue decomposition is the more informative measurement, because it says which protein supplies the binding enthalpy. In US WI-09, with POS left to locate its own site from 10 Å away, VP0 residues account for 86% of the receptor contribution; in all four resistant trajectories no VP0 residue enters the pocket at all, and the leading anchor is instead VP3 Trp24. Comparable enthalpy is therefore supplied by a different subunit, not less enthalpy by the same one. The genetics agree: VP0 Lys66 is the only position structurally linked to bound POS, and only in US WI-09, while Y224C mutation and the Arg224 that 9C-CMKC carries naturally both abolish the Tyr224-VP3 anchor. Resistance therefore arises either by perturbing the functional site (as at residue 66), or by relocating the ligand away from it (as at residue 224). Only the second route is open to a virus that has never seen the drug.

The infection data show whether this difference in binding site translates into difference in antiviral activity. At 0.3 µM, POS reduced both isolates in Vero-P cells but in MYADM-overexpressing cells suppressed only US WI-09; equal inhibition of US WI-09 in both contexts is consistent with restraint of a post-attachment VP0-dependent step rather than competition for attachment, while residual 9C-CMKC sensitivity in Vero-P plausibly reflects kinetic drag from peripheral VP3 occupancy at an already-limiting step. Vero-P, the condition in which POS was originally characterized, is thus precisely where a relocated, non-productive ligand still scores as a partial hit. Because the 2022 clade carries the VP1 224 substitution and the VP0 A124T allele as natural sequence, contemporary PeV-A3 arrives pre-adapted to evade the only compound class reported to inhibit it while still binding it avidly. POS should not advance without isolate-level stratification; VP1 224, read with VP0 66 and 124, is a companion marker available from specimens already sequenced for genotyping. Since PeV-A3 lacks the hydrophobic VP1 pocket that enterovirus capsid binders exploit, the VP0 hotspot defined here is the first functionally validated potentially druggable target on this capsid.

These conclusions are bounded in four ways. Two protomers probe interfacial chemistry rather than uncoating. A 500 ns trajectory cannot report an off-rate. The site assignments rest on reliable molecular dynamics and energetic convergence rather than static docking. And with one isolate per clade, our attribution of relocation to loss of the Tyr224 anchor stays inferential; isogenic 224 reversion and a cryo-EM POS complex are the decisive tests. The generalization is about method. Five choices did the work here. We co-folded across real clinical isolates alongside a genetically defined resistant control. We added a displaced-ligand rebinding control, so that the drug selected its own site. We defined pockets by occupancy across the ensemble rather than from single frames. We read per-residue enthalpy by subunit rather than in total. And we ran the infection assays in two receptor contexts. Together these convert “the compound binds the capsid” into a testable claim about which surface must be occupied for inhibition. The same gap recurs across *Picornaviridae*. Interprotomer pockets are assigned relevance from proximity, and virtual screens rank by affinity into pockets of unverified consequence. The workflow is therefore portable: to pleconaril- and vapendavir-class binders in EV-A71, EV-D68 and rhinovirus, and to the untested susceptibility of PeV-A1 and the emerging PeV-A5 lineages. Define the functional surface first, then use its occupancy as the primary screening endpoint. That is what turns capsid-directed antiviral work from affinity optimization into mechanism-guided drug discovery.

## ACKNOWLEDGMENTS

We would like to acknowledge collaborators at the Centers for Disease Control and Prevention, Atlanta, for providing Vero-p cell line, and Jan Carette Lab at Standford Medicine, USA, and Matthew Vogt Lab at University of North Carolina at Chapel Hill, USA for providing Vero-MYADM cell line. R.C. is grateful to NSF 22-599, EPSCoR RII Track-1, Award Number #2242763 for the computational resources used in this proposal.

## AUTHOR CONTRIBUTIONS

The study was conceived by R.C., R.S and D.D. Simulations and analyses were run by R.Sa. Experiments were run by D.D. under supervision of R.S. All authors contributed to the manuscript writing.

