## Supplementary for "Structural characterization of Posaconazole- parechovirus A3 capsid interaction using molecular simulations and cell infection assays"

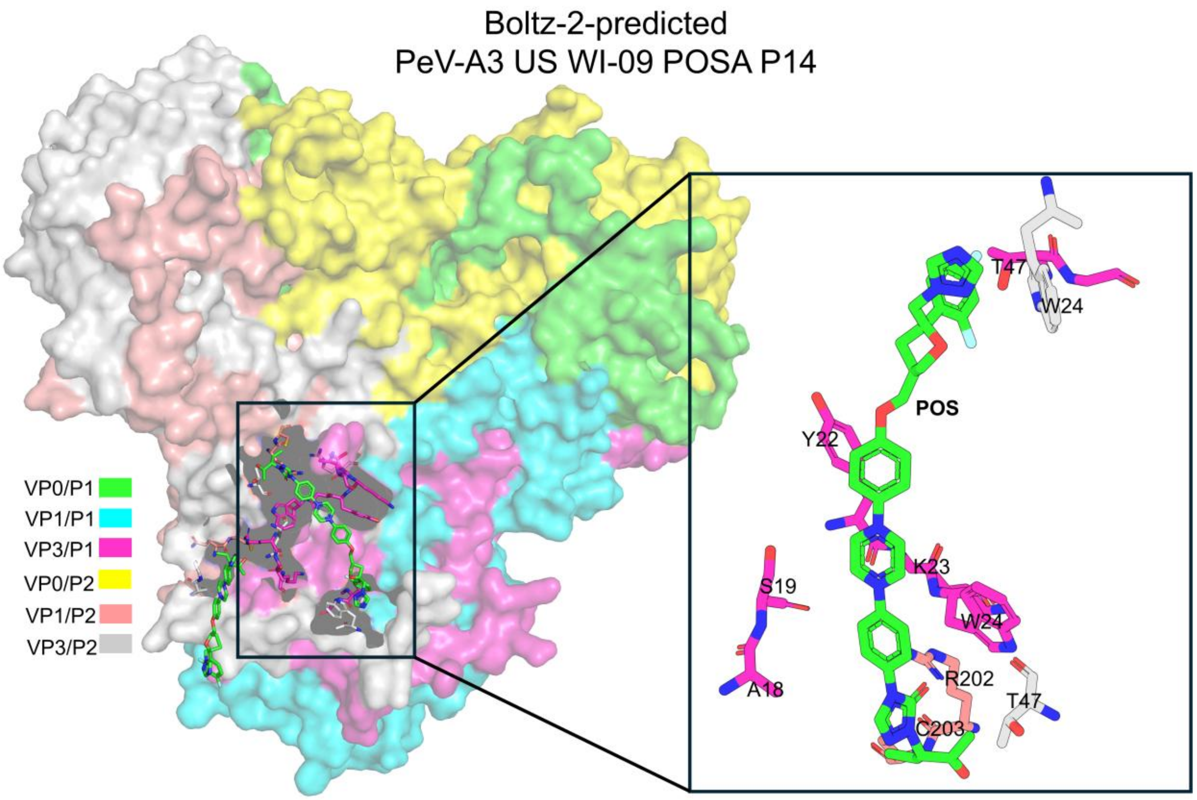


**SI Figure 1**. Boltz-2 two-protomer POS complex for US WI-09 POSA P14. POS binds the VP3 N-terminal arm (A18, S19, Y22, K23, W24) together with T47, with VP1 contacts at R202/C203, matching the VP3–VP1 site rather than the wild-type VP0–VP1 face.

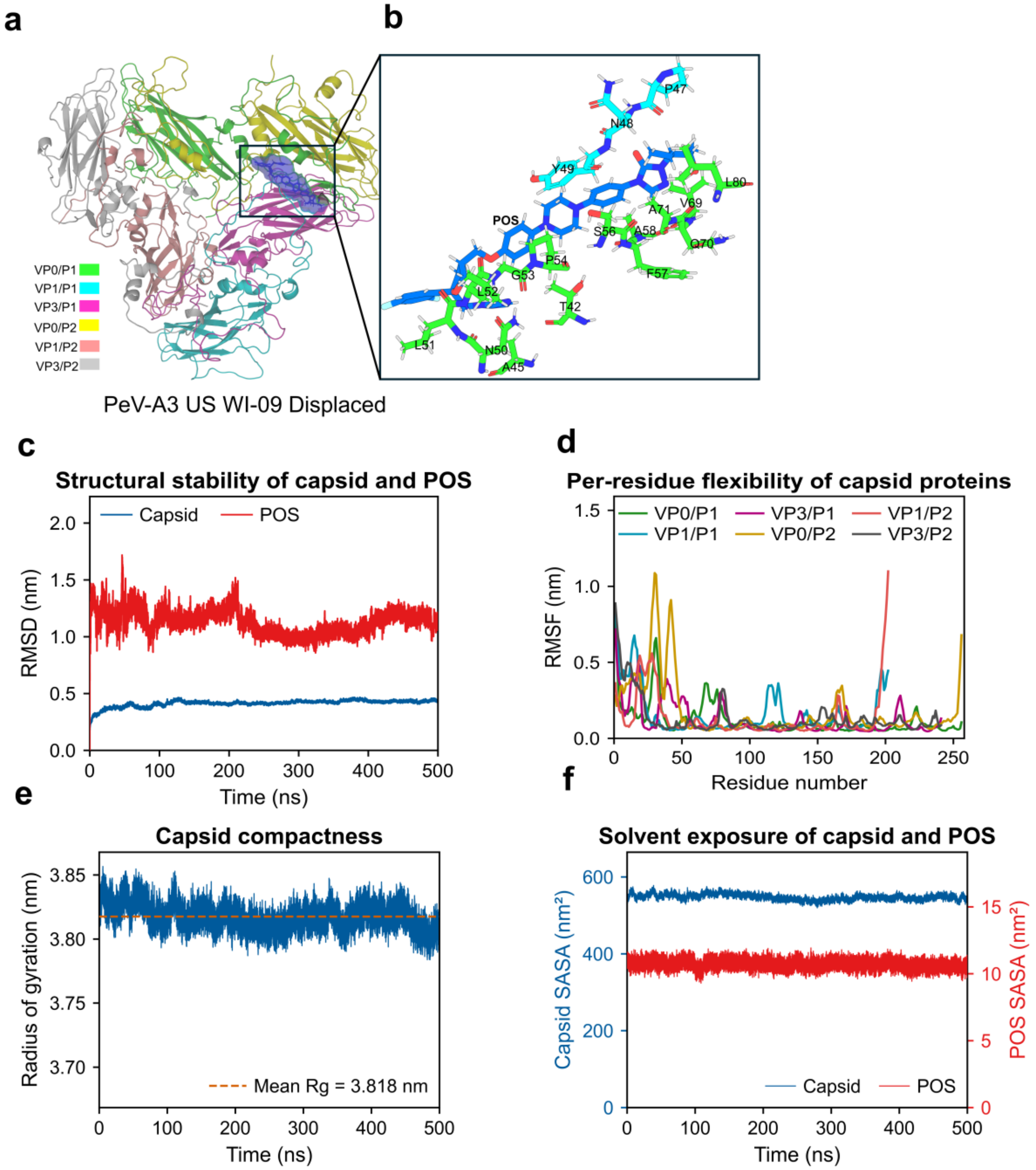


**SI Figure 2**. POS binding and stability in the displaced PeV-A3 US WI-09 system. a) Whole two-protomer structure at t = 500 ns, chains colored as in the legend, with POS shown as spheres. b) Close-up of the POS pocket on the VP0–VP1 face; after ~10 Å displacement POS returns to VP0 and recruits additional VP0/VP1 contacts (sticks, labeled). c) Backbone RMSD of the capsid and of POS over 500 ns. d) Per-residue Cα RMSF by chain. e) Capsid radius of gyration (mean Rg = 3.818 nm, dashed). f) Solvent-accessible surface area of the capsid and of POS.

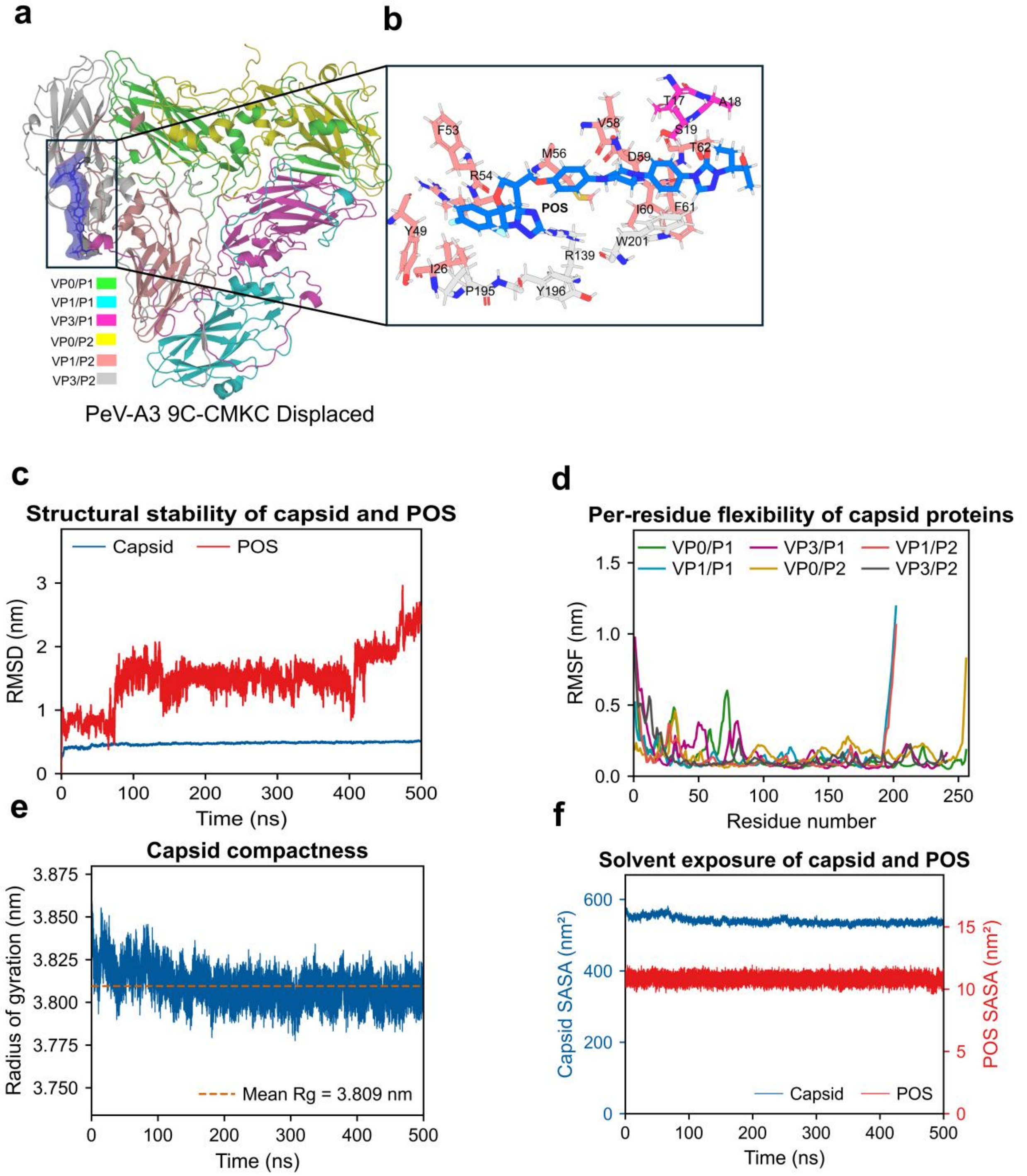


**SI Figure 3.** POS binding and stability in the displaced PeV-A3 9C-CMKC system. a) Two-protomer structure at t = 500 ns, chains colored as in the legend, with POS shown as spheres. b) Close-up of the final POS pocket; after ~10 Å displacement POS returns to the VP3–VP1 site and, over the last 100 ns, settles onto VP1 I60/T62 and VP3 W201 with the VP3 arm (W24) released. Contacting residues are shown as sticks and labeled. c) Backbone RMSD of the capsid and of POS. d) Per-residue Cα RMSF by chain. e) Capsid radius of gyration (mean Rg = 3.809 nm, dashed). f) Solvent-accessible surface area of the capsid and of POS.

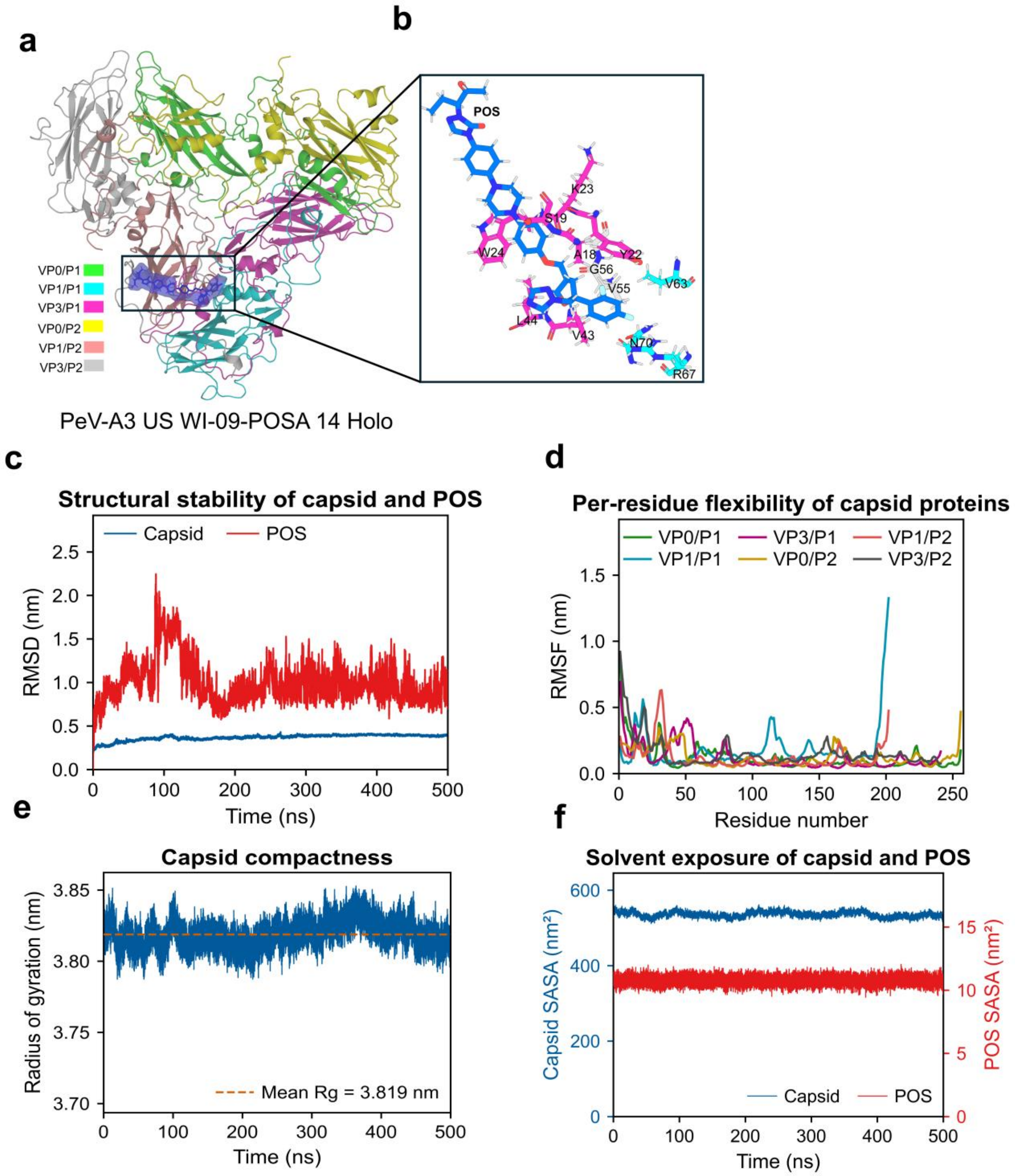


**SI Figure 4.** POS binding and stability in US WI-09 POSA P14 (holo). a) Two-protomer structure at t = 500 ns, chains colored as in the legend, with POS shown as spheres. b) Close-up of the POS pocket at the VP3 N-terminal annulus; the aromatic clamp (Y22, W24) and the inter-protomer hub V55 are shown with the other contacting residues as sticks and labeled. c) Backbone RMSD of the capsid and of POS over 500 ns. d) Per-residue Cα RMSF by chain. e) Capsid radius of gyration (mean Rg = 3.819 nm, dashed). f) Solvent-accessible surface area of the capsid and of POS.

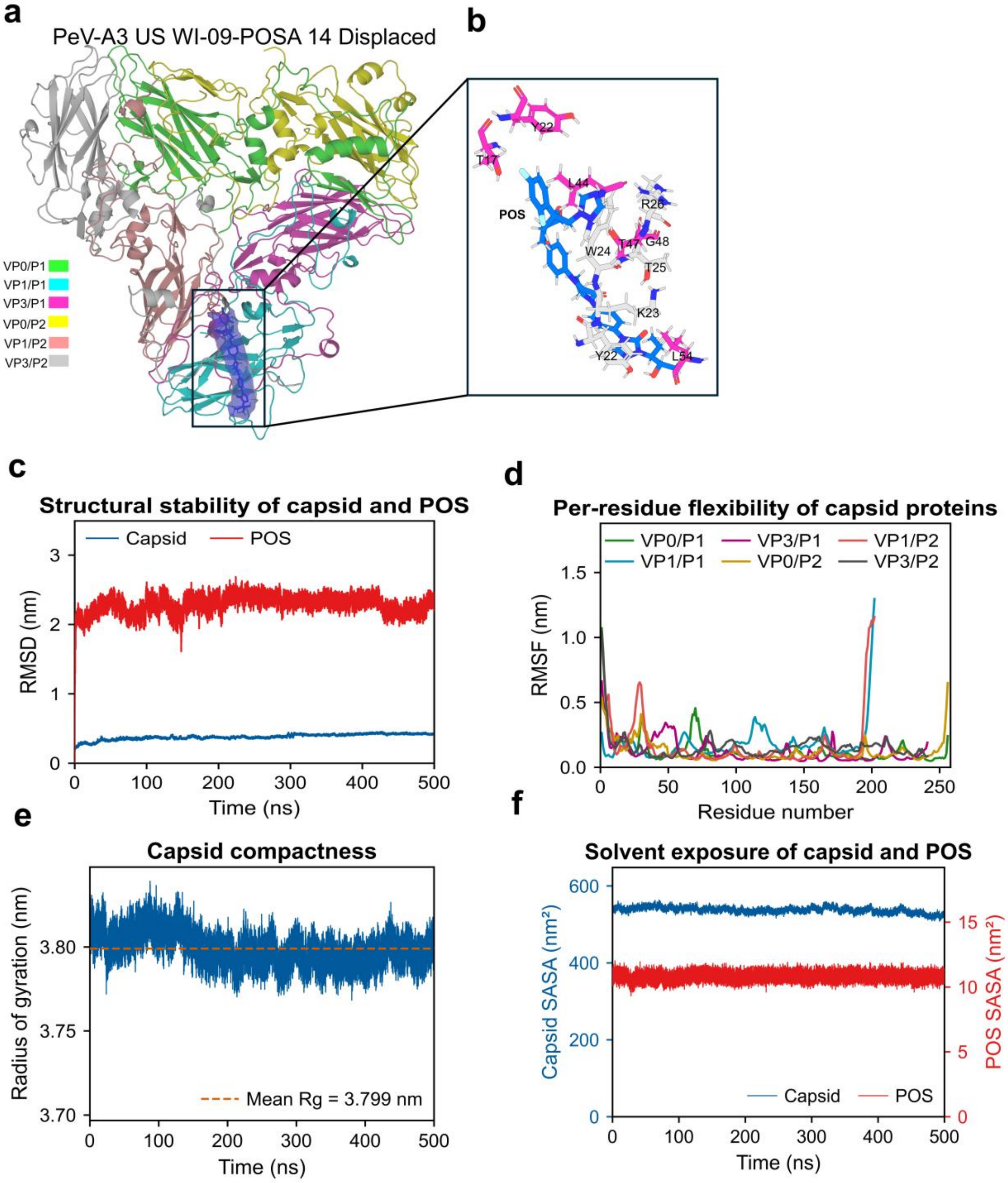


**SI Figure 5.** POS binding and stability in the displaced US WI-09 POSA P14 system. a) Two-protomer structure at t = 500 ns, chains colored as in the legend, with POS shown as spheres. b) Close-up of the final POS pocket; after ~10 Å displacement the Y22/W24 aromatic clamp re-forms on the opposite protomer (VP3 of protomer 2), identifying the two-fold VP3–VP3 arm cleft. Contacting residues are shown as sticks and labeled. c) Backbone RMSD of the capsid and of POS. d) Per-residue Cα RMSF by chain. e) Capsid radius of gyration (mean Rg = 3.799 nm, dashed). f) Solvent-accessible surface area of the capsid and of POS.

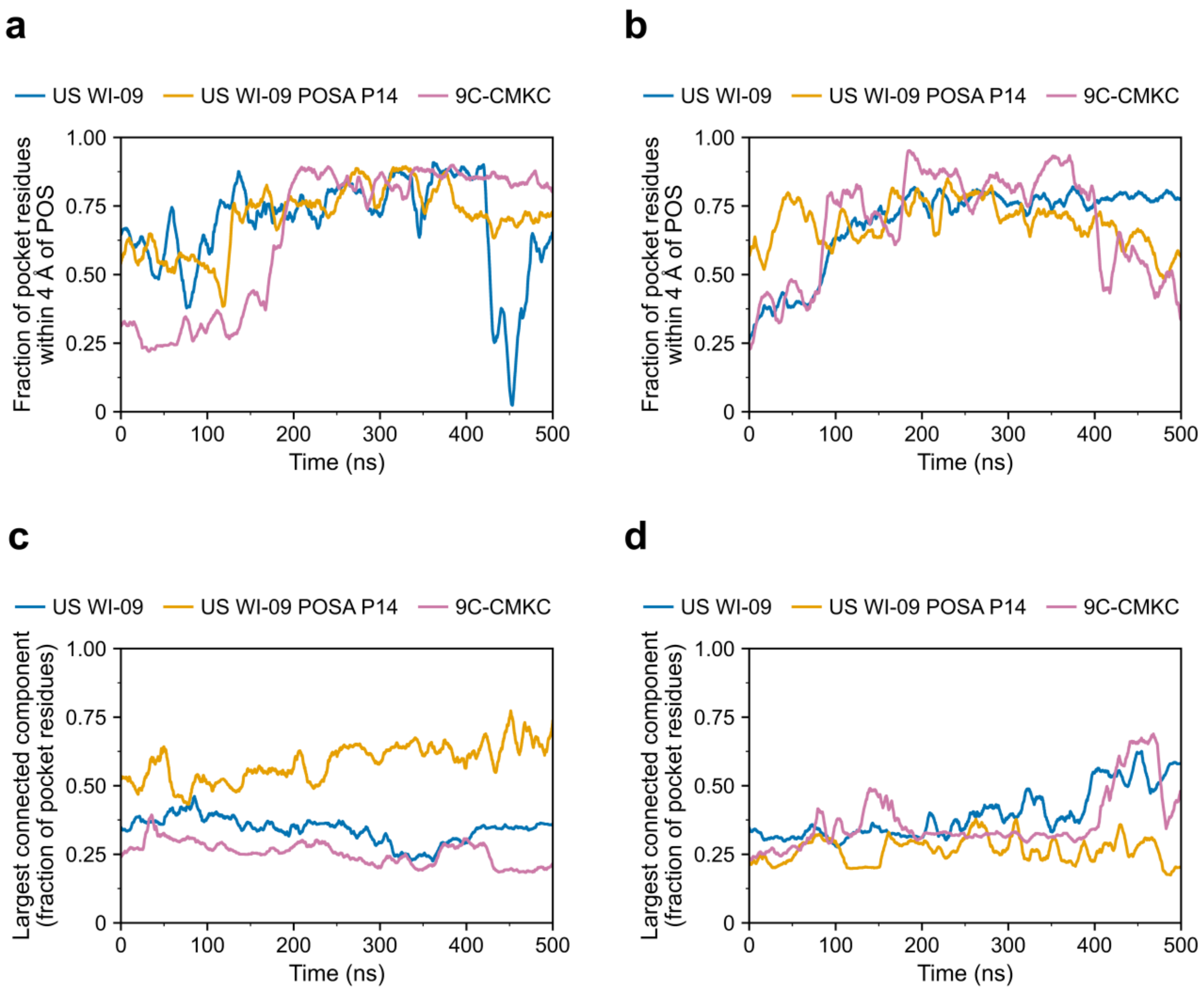


**SI Figure 6.** POS pocket engagement and cohesion over 500 ns for US WI-09, 9C-CMKC and US WI-09 POSA P14. Engagement is the fraction of pocket residues with any heavy atom within 4 Å of POS per frame; the largest connected component (LCC) is the fraction of pocket residues forming one connected cluster (side-chain centroids within 6 Å), a measure of how tightly the pocket holds together. a) Engagement, holo systems. b) Engagement, displaced systems. c) LCC, holo systems. d) LCC, displaced system.

**1. Computational binding affinity is not the ultimate determinant of site of action of POS.** Binding energetics were computed with gmx_MMPBSA(1) using the single-trajectory protocol of Miller et al. (2), over 1,000 frames sampled at 0.5 ns intervals from each 500 ns trajectory. The receptor was the occupancy-defined pocket, comprising residues within 4 Å of POS in at least half of all frames, so that energetic hotspots map onto the same contact set used in Sections 3.2 and 3.3, and per-residue decomposition was performed at the same resolution (idecomp = 2). Comparable workflows have been applied to viral capsid complexes after extended MD (1, 3). As pocket size differed among isolates, ΔH was normalised per residue for comparison.

Interaction entropy followed Duan et al. (4). gmx_MMPBSA treats IE as unreliable when the standard deviation of the interaction energy, σ_int, exceeds ~3.6 kcal/mol, consistent with the convergence criterion implemented in gmx_MMPBSA(5). That threshold is exceeded in all six systems (5.73 to 12.94 kcal/mol; SI Table 1). We therefore did not rank genotypes by −TΔS or by ΔG, and used the entropy-excluded binding enthalpy instead, as is standard when entropy estimates are poorly converged (6, 7). The consequence of ignoring this is severe: ranking the holo systems by ΔG would make 9C-CMKC the most favorable complex (−9.77 kcal/mol) and leave US WI-09 formally unbound (+7.76 kcal/mol), an ordering that reverses the cell data and reflects an unstable entropy term rather than the chemistry of the complexes. ΔH is favorable in all six ensembles, from −14.69 to −29.36 kcal/mol (SI Table 1). Once normalised for pocket size the three holo systems converge to −1.34, −1.75 and −1.36 kcal/mol per residue, so the 8 kcal/mol spread in total ΔH is largely a consequence of pocket composition. This agrees with the contact analysis, in which pocket engagement was statistically indistinguishable across all six runs ($|z| \leq 0.43$). Resistance does not arise from weaker binding, and ΔH does not identify the productive site. MM/PBSA is appropriate for relative comparisons among comparable receptor-ligand systems and is sensitive to the binding environment, so these energies should not be read as surrogates for antiviral efficacy (7). Capsid-directed antivirals act by engaging functionally important capsid structures (8) so favorable energetics at an alternative site are not evidence of preserved activity.

Per-residue decomposition resolves the split (SI Figure 7). In US WI-09 the enthalpy is spread thinly across the VP0-VP1 contact wall, with no residue exceeding one kcal/mol: VP1 Tyr49 (−0.96 ± 0.18), VP0 Leu51 (−0.95 ± 0.26), VP0 Gly53 (−0.80 ± 0.08) and VP1 Phe53 (−0.67 ± 0.21), with a ligand term of −8.65 ± 0.87 kcal/mol. That set reproduces the Phe/Tyr wall and VP0 packing described in Section 3.3. In the US WI-09 POSA P14 the enthalpy is larger and concentrated on a single VP3 aromatic anchor, Trp24 at −2.78 ± 0.21 kcal/mol, followed by Leu44 (−1.22 ± 0.03), Tyr22 (−1.01 ± 0.06) and Ala18 (−0.73 ± 0.18), with the most favorable ligand term of the three holo systems (−11.68 ± 1.05). 9C-CMKC occupies the same VP3 arm but distributes its energy across both protomers and both proteins, led by VP3 Phe60 (−1.87 ± 0.58), VP3 Trp24 (−1.22 ± 0.53), VP3 Leu44 (−1.12 ± 0.24) and VP1 Thr62 (−1.04 ± 0.36), with a ligand term of −10.54 ± 1.75. The decomposition assigns US WI-09 POSA P14 extra favorable contribution to specific VP3 aromatic contacts, not to any return of the VP0 surface.

VP0 is the single feature that separates US WI-09 from both 9C-CMKC and US WI-09 POSA P14. VP0 contributes five pocket residues and −1.31 kcal/mol in US WI-09 holo, and in the displaced run, where POS begins 10 Å away and locates its own site, sixteen VP0 residues supply −8.66 kcal/mol, 86% of the receptor total (SI Figure 8). In 9C-CMKC and US WI-09 POSA P14 trajectories VP0 contributes exactly zero: no VP0 residue enters the pocket in either the pre-bound or the displaced configuration, in either genotype. Both resistant backgrounds converge instead on the VP3 N-terminal arm from an unbiased start, with Trp24 the leading anchor in US WI-09 POSA P14 (−2.78 holo, −3.36 ± 0.08 displaced) and in 9C-CMKC displaced (−1.31 ± 0.41), and second only to Phe60 in 9C-CMKC holo. Convergence of the displaced runs onto the sites found in the holo runs indicates genuine energetic minima rather than artefacts of the docked pose and shows that VP0 exclusion is a property of the resistant capsid rather than of the starting structure.

The US WI-09 VP0 site and the US WI-09 POSA P14 VP3 site differ in chemistry. Aromatic residues account for 41% of the receptor enthalpy in US WI-09 holo and 15% in US WI-09 displaced, where the site is a shallow groove dominated by aliphatic packing (VP0 Pro54 −1.90 ± 0.22, VP1 Tyr49 −1.40 ± 0.29, VP0 Val69 −1.28 ± 0.28). In the resistant systems aromatics supply 50 to 70%, reaching an extreme in the displaced US WI-09 POSA P14, where Trp24 and Tyr22 of a single protomer account for −5.33 of −8.26 kcal/mol. Binding is driven throughout by shape complementarity rather than electrostatics (ΔE_vdW −27.06 to −45.97 against ΔE_ele −3.91 to −10.65 kcal/mol), with polar desolvation offsetting 57 to 75% of the van der Waals gain, which is why no residue contributes more than 3.4 kcal/mol even at the strongest anchor.

Enthalpy therefore reports how well POS packs into whichever site it occupies, while antiviral outcome depends on whether that site lies on the entry-relevant VP0 surface. US WI-09 pays its binding energy on VP0 and VP1 in both starting conditions. Both resistant genotypes pay a comparable or larger per-residue enthalpy on a VP3 aromatic pocket while leaving VP0 entirely ligand-free. Enthalpically concentrated packing at the non-productive VP3 site in US WI-09 POSA P14 and more dispersed packing at the same site in 9C-CMKC produce the same functional outcome: POS does not occupy the VP0 surface. VP0 has been identified experimentally as the PeV-A3 capsid protein that engages the essential entry factor MYADM (9), which gives a structural basis for loss of antiviral activity without any loss of binding affinity.

**SI Table 1**. MM/PBSA binding energetics for the holo and displaced POS-capsid complexes (kcal/mol; 1,000 frames at 0.5 ns intervals; interaction entropy over the last 250 frames). ΔH, entropy-excluded binding enthalpy; −TΔS, interaction-entropy term; ΔG = ΔH + (−TΔS), reported as mean ± block SEM; σ_int, standard deviation of the interaction energy.

| **System** | | **ΔH (kcal/mol)** | **-TΔS (kcal/mol)** | **ΔG (kcal/mol)** | **σ_int (kcal/mol** |
|---|---|---|---|---|---|
| US WI-09 | Holo | −14.69 | +22.45 | +7.76 ± 8.84 | 9.28 |
| | Displaced | −29.36 | +10.53 | −18.83 ± 7.55 | 11.58 |
| US WI-09 POSA P14 | Holo | −22.81 | +17.83 | −4.98 ± 5.71 | 10.43 |
| | Displaced | −21.99 | +12.53 | −9.46 ± 3.39 | 5.73 |
| 9C-CMKC | Holo | −21.80 | +12.03 | −9.77 ± 8.57 | 12.94 |
| | Displaced | −21.99 | +19.47 | −2.53 ± 8.59 | 12.10 |

†ΔG= ΔH + (−TΔS). Not used for ranking because σ_int > 3.6 kcal/mol in all rows

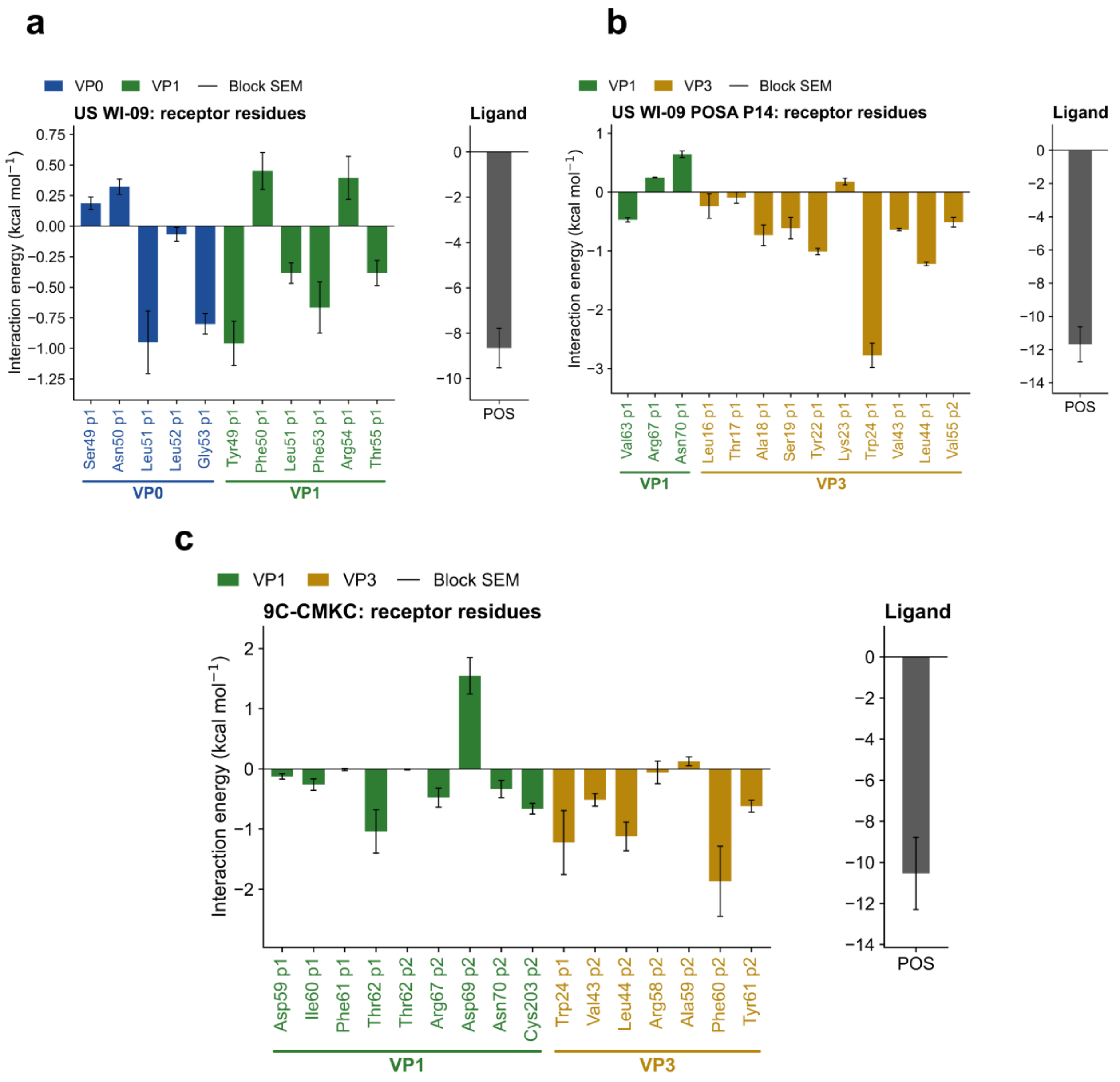


**SI Fig 7.** Per-residue MM/PBSA decomposition of the holo complexes. a) US WI-09, b) US WI-09 POSA P14, c) 9C-CMKC. Bars give each receptor residue's net contribution to the binding enthalpy, colored by capsid protein (VP0, VP1, VP3), with the POS self-term shown in the adjacent panel. Values are mean ± SEM across five consecutive 100 ns blocks; treating frames as independent would understate the error roughly tenfold. VP0 contributes to US WI-09 only; both resistant genotypes draw their enthalpy from the VP3 N-terminal arm.

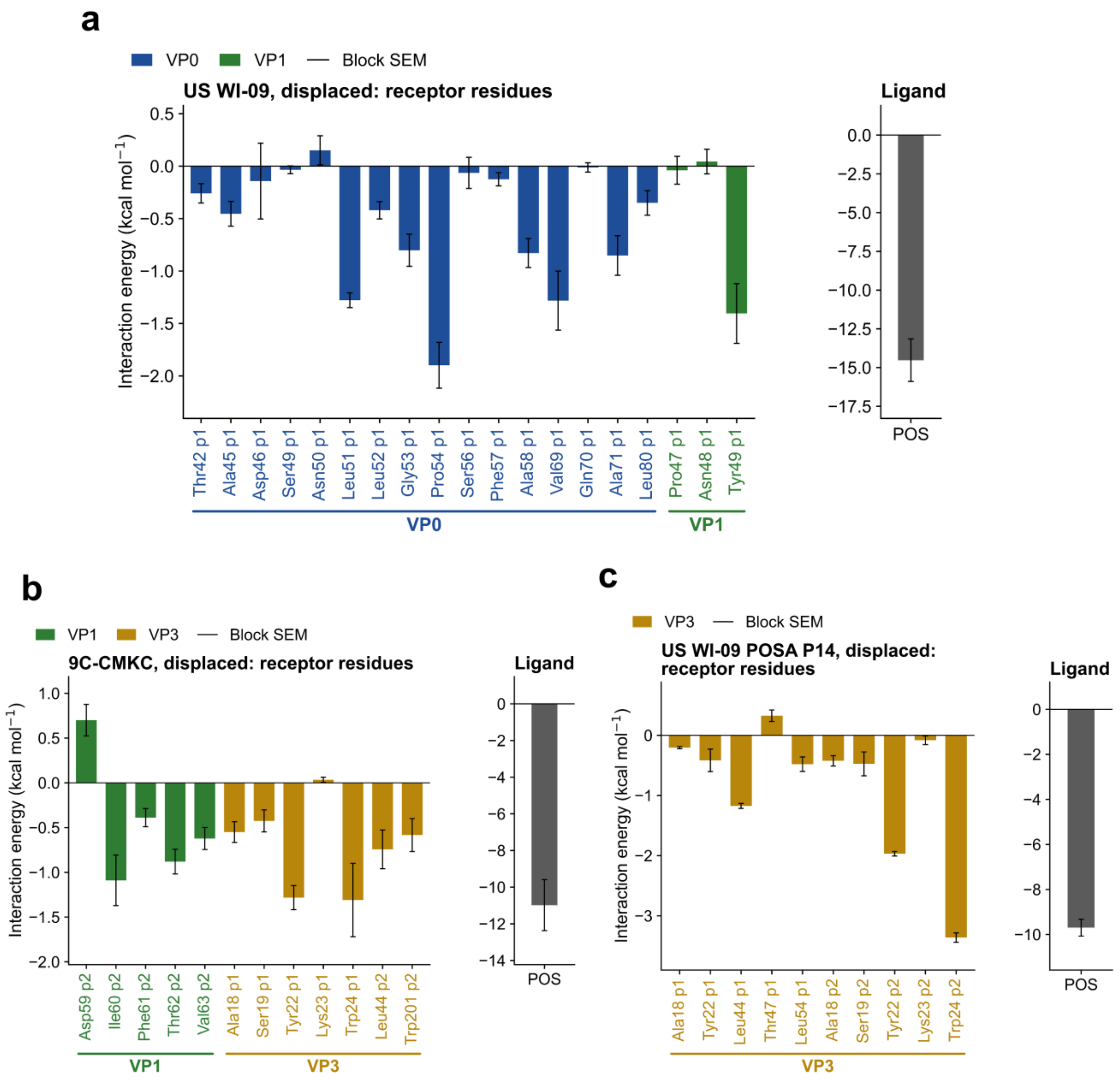


**SI Fig 8.** Per-residue MM/PBSA decomposition of the displaced complexes. a) US WI-09, b) 9C-CMKC, c) US WI-09 POSA P14. After ~10 Å displacement, US WI-09 recovers a VP0-dominated site (16 VP0 residues, 86% of the receptor enthalpy), while both resistant genotypes again place all enthalpy on VP3/VP1 with no VP0 contribution, confirming VP0 exclusion is a property of the resistant capsid rather than the starting pose.
